# Treatment-free interval duration during T-cell engager conditioning shapes simulated T-cell exhaustion and post-transfer tumor control

**DOI:** 10.1101/2025.11.17.688873

**Authors:** Nina Obertopp, Matthew Froid, Shari Pilon-Thomas, David Basanta

**Affiliations:** Cancer Biology Ph.D. Program, University of South Florida, Tampa, Florida, USA; Department of Immunology, H. Lee Moffitt Cancer Center & Research Institute, Tampa, Florida, USA; Department of Integrated Mathematical Oncology, H. Lee Moffitt Cancer Center & Research Institute, Tampa, Florida, USA

## Abstract

**Background:** Blinatumomab, a CD3/CD19 bispecific T-cell engager (TCE), is effective in relapsed/refractory (R/R) B-cell acute lymphoblastic leukemia (B-ALL), but responses may be limited by T cell exhaustion. A recent preclinical study found that incorporating a 7-day treatment-free interval (TFI) into TCE exposure reduced T cell exhaustion and improved subsequent antitumor activity compared with continuous exposure.

**Methods:** Based on this preclinical study, we developed an agent-based model (ABM) to evaluate alternative TFIs by simulating T cell–tumor cell interactions, in which TCE availability influences T cell activation, tumor-cell killing, and T cell exhaustion. To reproduce the experimental design, simulations included a 14-day in vitro conditioning phase with varying TCE schedules, followed by an adoptive cell transfer (ACT) phase in which post-transfer tumor control was evaluated.

**Results:** Our ABM reproduced the experimentally observed reduction in CD8⁺ T cell exhaustion following a 7-day TFI versus continuous TCE exposure (CONT). Among tested TFIs, the 7-day TFI produced the lowest exhaustion (p_adj = 3.52 × 10⁻¹⁷ vs. CONT) and lowest tumor burden (p_adj = 1.93 × 10⁻¹⁶ vs. CONT). The 5-day TFI was the only other tested schedule to significantly reduce tumor burden relative to CONT. Shifting the 7-day TFI from the second to the first week of conditioning altered T cell phenotype at harvest but not tumor control during the ACT phase (p_adj = 0.87). Limiting TCE exposure during the in vitro phase to one day further reduced tumor burden (all p_adj < 5 × 10⁻¹⁷), with exposure on the first day outperforming exposure immediately before T cell harvest (p_adj = 3.6 × 10⁻¹⁰).

**Conclusions:** The benefit of TFIs depends on interval duration and schedule organization. Our findings also highlight a potential limitation of current preclinical TFI testing. Because tumor cells are replenished rather than allowed to grow during in vitro conditioning, prolonged TFIs carry no penalty from tumor growth. Models that apply TFIs directly to growing tumors are needed to determine whether reduced T cell exhaustion outweighs tumor progression during treatment withdrawal.

**Author Summary:** T cell engagers help T cells recognize and attack tumor cells. Although these treatments can help control tumors, they may also overstimulate T cells, causing them to become exhausted and lose function. Temporarily interrupting treatment could give T cells time to recover, but testing the many possible ways to schedule these treatment breaks experimentally would be difficult. We therefore developed a computer model that simulates T cells and tumor cells to investigate how the duration and timing of treatment breaks influence T cell exhaustion and subsequent tumor control. Our model reproduced experimental findings showing that treatment breaks can preserve T cell function. We then used the model to compare additional schedules. In general, longer treatment breaks preserved T cell function better than several shorter breaks. We also examined whether the placement of the break within the treatment schedule mattered and found that moving the same break to an earlier point did not affect subsequent tumor control. Our findings show how computer models can help compare treatment schedules and reveal that more realistic models and experimental testing are needed.

## Introduction

B-cell acute lymphoblastic leukemia (B-ALL) is a malignancy of immature B-cell precursors that arises in the bone marrow and disrupts normal hematopoiesis. While frontline chemotherapy induces long-term remission in approximately 80–85% of children and 40–50% of adults with newly diagnosed B-ALL, outcomes remain poor for patients with relapsed or refractory (R/R) disease.^1^ Recent advances in T cell-based immunotherapies, including chimeric antigen receptor (CAR)-T cell therapy and bispecific T cell engagers (TCEs), have significantly expanded treatment options for patients with R/R B-ALL. Unlike CAR-T cell therapy, which requires ex vivo engineering and expansion of autologous T cells, TCEs are administered as off-the-shelf agents capable of rapidly redirecting endogenous T cells toward malignant cells. The CD3/CD19-directed bispecific TCE blinatumomab, FDA-approved since 2014, has shown substantial activity in both adult and pediatric patients with R/R B-ALL, inducing complete remission in up to 69% of patients.^2^ While blinatumomab has reshaped treatment options and is increasingly incorporated into frontline regimens, resistance, relapse, and modest overall survival highlight the need to better understand and overcome mechanisms of treatment failure.^1^

As with many immunotherapies, treatment failure during TCE therapy is frequently associated with increased T cell exhaustion.^3^ Applying treatment-free intervals (TFIs) to interrupt continuous TCE administration (CONT) may provide an opportunity for T cells to recover functional capacity. This concept is supported by observations in CAR-T cell therapy, where transient pharmacologic interruption of chronic CAR signaling using dasatinib restored effector functionality and remodeled exhaustion-associated transcriptional programs.^4^ Treatment-free periods are already incorporated into the clinical administration of blinatumomab, although not specifically for the purpose of mitigating T cell exhaustion. Adults with R/R B-ALL typically receive 42-day treatment cycles consisting of 28 days of continuous intravenous infusion followed by a 14-day treatment-free period.^6^ This treatment-free period was originally included in the trial schema as a safety window for adverse-event monitoring and infusion logistics, rather than for an anticipated biological benefit, and was retained in labeling after regulators observed that peripheral B-cell suppression persisted during the break.^7^ To test whether an interruption within the initial 28-day exposure period could provide an additional biological benefit, Philipp et al. incorporated a 7-day TFI into an in vitro TCE model. This reduced T cell exhaustion and generated T cells with greater antitumor activity following adoptive cell transfer (ACT) into tumor-bearing mice.^5^

While Philipp et al. showed that a 7-day TFI improved T cell function, this schedule was selected a priori rather than through systematic evaluation of alternative schedules.^5^ We therefore asked how the duration and timing of TFIs influence T cell phenotype and subsequent antitumor activity, and whether the benefits of TFIs are determined primarily by the overall reduction in TCE exposure or by how that exposure is temporally organized. To address these questions, we calibrated an agent-based model (ABM) using the experimental data reported by Philipp et al.^5^ and systematically evaluated alternative TFI schedules.

ABMs have been widely used in computational cancer modeling to investigate T cell exhaustion, immunotherapy, resistance mechanisms, and treatment scheduling.^8–10^ Unlike population-based modeling approaches, ABMs simulate individual cell behaviors and interactions, allowing heterogeneous T cell states and their dynamic responses to treatment exposure and withdrawal to emerge from defined biological rules.^11^ Here, we developed an ABM to investigate T cell activation, proliferation, exhaustion, and cytotoxicity during TCE exposure while systematically varying the duration and timing of TFIs. We then evaluated how these schedules shaped the resulting T cell phenotype and its antitumor activity following ACT. This approach enables broad comparison of TFI schedules that would be impractical to evaluate exhaustively in preclinical models.

## Methods

Mathematical modeling has long been recognized as a powerful and efficient approach for testing and predicting treatment strategies in cancer.^12^ Among these approaches, ABMs represent a particularly versatile class of computational models in which individual entities, or agents, follow predefined behavioral rules and interact with each other and their environment. Building on foundational single-cell models of in vitro spatial dynamics,^13^ modern ABMs leverage diverse open-source and hybrid modeling frameworks^14,15^ to capture the highly localized, multi-scale nature of the tumor immune microenvironment.^16^ By representing local interactions among individual cells, ABMs allow complex population-level behaviors to emerge from relatively simple agent-level rules. This bottom-up, stochastic approach is now well established in cancer research and is widely used to represent biological complexity.^17^

### Agent-based model overview

Our ABM was developed as a Hybrid Cellular Automata (HCA), which has been widely applied to investigate cellular interactions and evolutionary dynamics in diverse biological systems.^18^ The model was implemented using the Hybrid Automata Library (HAL).^19^ Discrete cellular agents follow predefined behavioral rules within a spatially explicit environment (Fig. 1). Each agent represents an individual tumor cell or CD8⁺ T cell. Because the model was calibrated to the CD8⁺ T cell exhaustion data reported by Philipp et al.,^5^ “T cells” refers specifically to modeled CD8⁺ T cells when describing the model. Agents reside on a two-dimensional lattice (160 × 150 sites) with periodic boundary conditions, with each lattice site accommodating at most one cell. Each time step corresponds to one hour, and agents are updated in randomized order at each step to prevent systematic bias arising from the update sequence. Local cell–cell interactions occur within a Moore neighborhood comprising the eight surrounding lattice sites and govern cell movement, T cell–mediated tumor killing, and tumor cell proliferation. Each T cell makes up to three movement or interaction attempts per step. Movement was not independently probabilistic beyond site availability, as an attempt succeeded whenever at least one unoccupied site was present in the Moore neighborhood, with one such site selected uniformly at random as the destination. The same selection method was used to place daughter cells following division. If no unoccupied site is available, the attempt is skipped.

**Figure 1.**
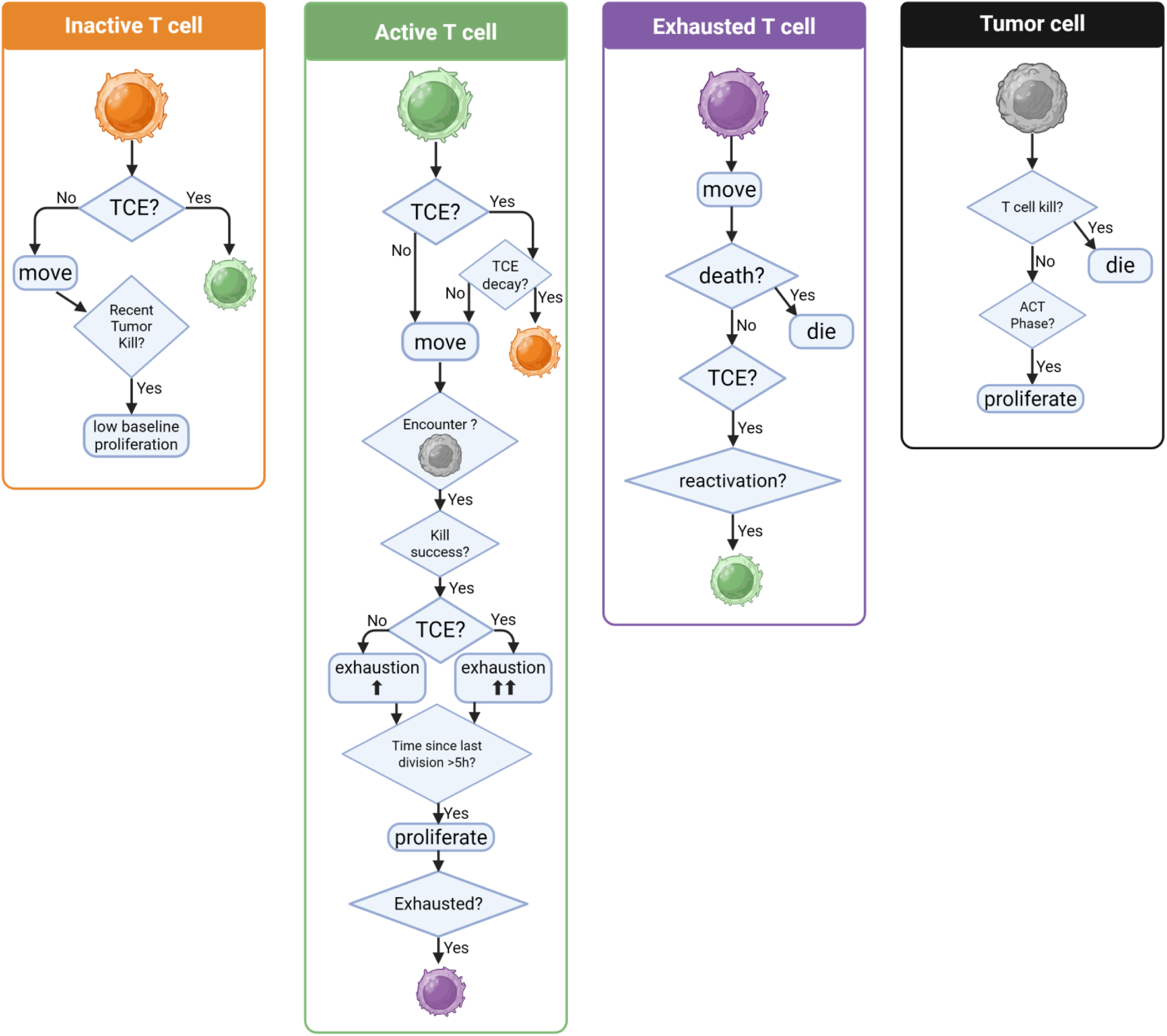
Agent-Based Model (ABM) Decision Tree of T Cell Engager (TCE) Therapy. Flowcharts show the decision logic governing each cell state at every simulation time step: inactive T cells (orange), active T cells (green), exhausted T cells (purple), and tumor cells (black). All modeled T cells represent CD8⁺ T cells. Inactive T cells move stochastically and become active when TCE is present, with recent tumor killing allowing low-probability basal proliferation. Active T cells move, engage and kill tumor cells upon encounter, accumulate exhaustion at a rate dependent on whether TCE is present, and proliferate if sufficient time has elapsed since their last division. Their probability of transitioning to the exhausted state increases with accumulated exhaustion. In the absence of TCE, active T cells revert to the inactive state. Exhausted T cells retain motility but do not kill or proliferate; they may die or rarely reactivate when TCE is present. Tumor cells proliferate during the adoptive cell transfer (ACT) phase and are removed following successful T cell-mediated killing.

### Model Initialization and Agent Rules

Our ABM reproduces the two-stage experimental design described by Philipp et al., which consisted of a 14-day in vitro conditioning phase followed by an ACT phase in which conditioned T cells were transferred into a patient-derived xenograft model of ALL (PDX-ALL-265).^5^

In accordance with the experimental design, the in vitro conditioning grid was initialized at 50% occupancy and an effector-to-target (E:T) ratio of 1:4, corresponding to 2,400 T cells and 9,600 tumor cells on the 24,000-site lattice. All T cells were initialized as inactive, representing the assumed baseline state of T cells before TCE-mediated activation. Tumor cells were modeled as non-proliferative to reflect the effects of irradiation before co-culture. Tumor cells were replenished every 7 days to restore an E:T ratio of 1:4 relative to the current T cell population, matching both the periodic addition of fresh target cells and the E:T ratio applied by Phillip et al.^5^ TCE exposure followed the schedule assigned to each simulated condition, including continuous exposure and the respective TFI schedules. Grid initialization and tumor cell replenishment used the same random placement method. Candidate lattice coordinates were sampled uniformly at random and re-sampled until an unoccupied site was found. Replenishment was implemented by removing all tumor cells present at the reset time point and reseeding new tumor cells at a 1:4 ratio relative to the current T cell count. When the current T cell population left fewer empty sites than required to reach this ratio, the number of tumor cells reseeded was capped to the number of available empty sites.

As described by Philipp et al.^5^ and reproduced in the ABM, simulated T cells were harvested at the end of conditioning on day 14 and transferred to a newly initialized lattice containing 1,000 PDX-ALL-265 tumor cells. As specified in the corresponding figure, either 200 T cells were randomly sampled, or all T cells from the in vitro phase were transferred; each cell was eligible for transfer at most once. Transferred T cells retained their functional state (inactive, active, or exhausted), accumulated exhaustion level (PD-1), and the time since their most recent tumor-cell kill (h). The ACT phase was then simulated for 16 days. Before transfer, an empty lattice was seeded with fresh tumor cells, after which the transferred T cells were placed using the random coordinate-sampling method described above.

### Tumor Cell Behavior

During the initial in vitro phase, tumor cells represent irradiated CD19⁺ OCI-Ly1 lymphoma cells and were therefore modeled as non-motile and non-proliferative. During the ACT phase, tumor cells represented PDX-ALL-265 leukemia cells and proliferated with a per-hour probability of 1/72. This value was calculated from pretreatment bioluminescence measurements reported by Philipp et al.^5^, which indicated an approximately 100-fold increase in tumor burden between days 14 and 28 after engraftment, before T cell or AMG 562 administration. Applying an exponential growth model to this increase yielded the proliferation probability without iterative calibration. In both phases, tumor cells were removed following a successful T cell–mediated killing event.

### T cell Behavior

Each T cell exists in one of three functional states: inactive, active, or exhausted. Each cell also carries a numerical exhaustion score, denoted by *E*, that governs transitions between functional states. Consistent with the use of peripheral blood T cells from healthy donors in the experimental study, all simulated T cells were initialized in the inactive state.^5^ This assumption is supported by the low expression of exhaustion-associated markers typically observed in peripheral blood T cells from healthy individuals.^26,27^

T cell exhaustion was modeled based on the experimental measurements reported by Philipp et al.^5^ In that study, exhaustion was quantified as the percentage of CD8⁺ T cells co-expressing the inhibitory receptors PD-1, TIM-3, and LAG-3. In the ABM, the cumulative exhaustion variable is labeled “PD-1” and denoted by *E*. Despite this label, *E* represents the combined exhaustion phenotype rather than the expression of PD-1 alone. Model calibration was performed against the experimentally measured percentages of exhausted CD8⁺ T cells under continuous and 7-day TFI schedules at the corresponding experimental time points.

Inactive T cells represent cells not actively engaged in TCE-mediated tumor killing. In the presence of TCE, inactive T cells transition to the active state. In the absence of TCE, inactive T cells attempt movement up to three times per hour, each attempt moving to an empty site drawn uniformly at random from the Moore neighborhood; an attempt with no available site was skipped. Once per hour, inactive T cells undergo low-frequency basal proliferation with a probability of 1/120, provided that a tumor-cell kill had occurred within the preceding 3 days. This represents antigen-nonspecific bystander proliferation supported by recent immune activity, consistent with the expansion of CD8⁺ T cells reported following IL-2-based combination immunotherapy.^22^ The daughter cell was placed at an empty neighboring site selected using the same method as for movement.

Active T cells migrate across the lattice and interact with adjacent tumor cells. Each active T cell attempts up to three movement or kill interactions per hour. During each interaction attempt, occupied sites in the Moore neighborhood were scanned sequentially. The first tumor cell encountered was selected for a killing attempt, provided at least 12 hours had elapsed since the T cell’s previous successful kill. This cooldown represents the finite serial-killing capacity of individual cytotoxic T cells. If it had not yet elapsed, the T cell instead attempted movement according to the same rules as an inactive T cell. Kill probability was computed as a sigmoidal function of the exhaustion score *E*:

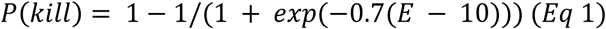

This function corresponds to 50% suppression at *E* = 10 and a steepness parameter of 0.7, such that the probability of successful killing decreases as *E* increases.^20,28^ This probability was doubled during the ACT phase compared with the in vitro phase, reflecting the greater susceptibility of PDX-ALL-265 cells used in the ACT experiments to TCE-mediated lysis relative to OCI-Ly1 cells used during in vitro conditioning, as reported by Philipp et al.^5^

Following each successful tumor-cell kill, *E* increases by a baseline increment of 1.0., and the active T cell becomes eligible for proliferation once at least 5 hours have elapsed since its previous division.^21^ Daughter T cells inherit the parental value of *E*, representing persistence of the acquired exhaustion phenotype during proliferation. This modeling assumption is consistent with evidence that exhaustion-associated transcriptional and epigenetic programs can remain stable during proliferative reinvigoration.^24^ Post-kill proliferation probability was set higher during the ACT phase compared to in vitro conditioning. This phase-specific difference was introduced to represent the substantial T-cell expansion observed following adoptive transfer in the experimental system,^5^ consistent with reports of rapid in vivo expansion of tumor-reactive T cells following ACT.^29^

The transition from the active to the exhausted state was modeled probabilistically as a sigmoidal function of the exhaustion score *E*:

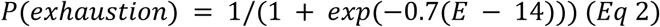

This function corresponds to a 50% exhaustion probability of transitioning to the exhausted state at *E* = 14 and a steepness parameter of 0.7. The probability of transitioning increases with (E), with the steepness parameter controlling the rate of this transition. This transition probability is evaluated once per movement or kill attempt (up to three times per hour) and compared with a uniform random draw to determine whether transition occurs.

Exhausted T cells retain motility but are modeled as noncytotoxic and non-proliferative. They are assigned a probabilistic death rate of 1/72 h⁻¹ to represent the reduced persistence of dysfunctional T cells, consistent with the evidence that prolonged stimulation can induce CD8⁺ T cell dysfunction together with activation-induced cell death.^23^ Exhausted T cells are modeled as largely dysfunctional but not irreversibly terminal. In the presence of TCE, exhausted T cells are assigned a low probability of transitioning back to the active state, representing rare partial functional reinvigoration consistent with experimental evidence that interruption and subsequent re-exposure to bispecific stimulation can restore effector function in a subset of T cells.^4^ Upon reactivation, 50% of *E* is retained as a modeling assumption, reflecting evidence that reinvigorated T cells can retain epigenetic features that limit durable reprogramming.^24^ The exhausted T cell death rate and reactivation probability were manually calibrated to the exhaustion dynamics reported by Philipp et al.^5^

### Modeling T cell engager (TCE) therapy and treatment-free intervals (TFI)

TCE exposure is represented as a spatially uniform binary variable indicating whether treatment was present or absent at each simulation time point. The model was parameterized to AMG 562, the half-life–extended CD19×CD3 bispecific antibody used in the experimental calibration dataset.^5^ AMG 562 has an approximately 210-hour half-life, substantially longer than the approximately 2-hour half-life of blinatumomab.^25^ The pharmacokinetic effect of AMG 562 withdrawal is incorporated into T cell state transitions, instead of explicitly modeling spatial or temporal drug concentrations. Following TCE withdrawal, active T cells retained their capacity to kill tumor cells while remaining in the active state. Active T cells subsequently revert to the inactive state according to an exponential decay function based on time since TCE withdrawal.

Tumor cell killing occurring while TCE was present increased *E* at a higher rate than kills occurring in its absence, accelerating progression from the active to the exhausted state. This reflects the mechanism of TCE-mediated activation, in which simultaneous engagement of CD3 on T cells and CD19 on tumor cells redirects T cells toward target cells independently of physiological T cell receptor (TCR) –major histocompatibility complex (MHC) recognition.^30^ Sustained TCE-mediated stimulation can promote rapid antitumor activity but also accelerate T cell exhaustion.^5,31^ This trade-off has motivated recent TCE design strategies aimed at reducing CD3-binding affinity while preserving antitumor activity.^32^

### Model calibration and sensitivity analysis

Model parameters and their corresponding baseline values and sources are summarized in Table 1. Calibration was based on the T cell exhaustion percentages reported by Philipp et al.^5^ for the CONT and 7-day TFI schedules at matched experimental timepoints. For parameters without a directly reported experimental value, including those with a literature-informed starting point, candidate parameter values were iteratively refined to improve agreement between simulated outcomes and the experimental calibration targets. These included the T cell kill and division cooldowns, proliferation rates, parameters governing kill suppression and exhaustion, exhausted T cell death, and reactivation probability (Table 1). No automated optimization algorithm was used. Calibration error was assessed using root mean squared error (RMSE), calculated from the deviations between simulated values and the corresponding experimental ranges. A deviation of zero was assigned when a simulated value fell within the reported range. The parameter set yielding the lowest RMSE relative to the target ranges was retained as the final calibrated set.

**Table 1.**
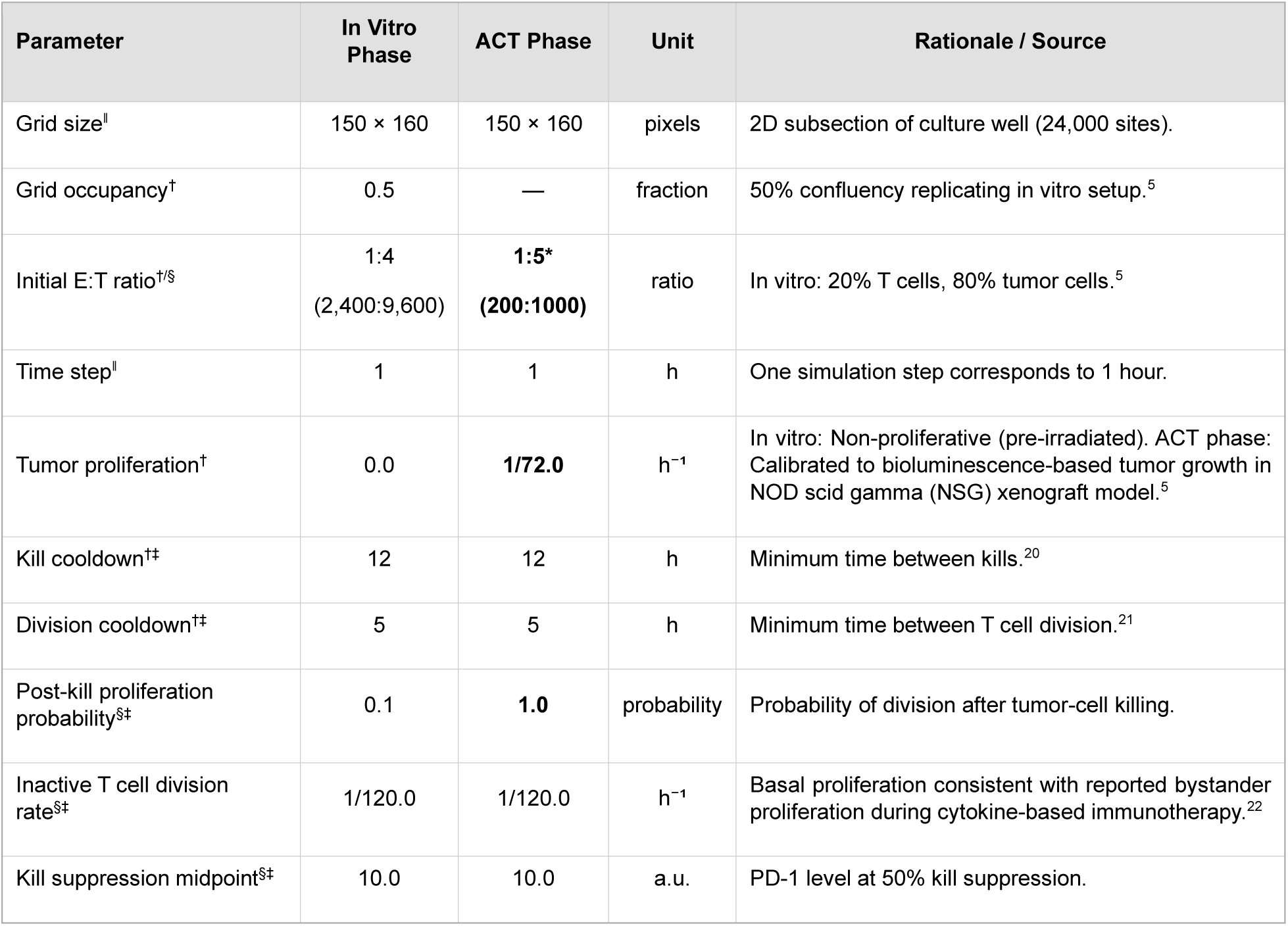
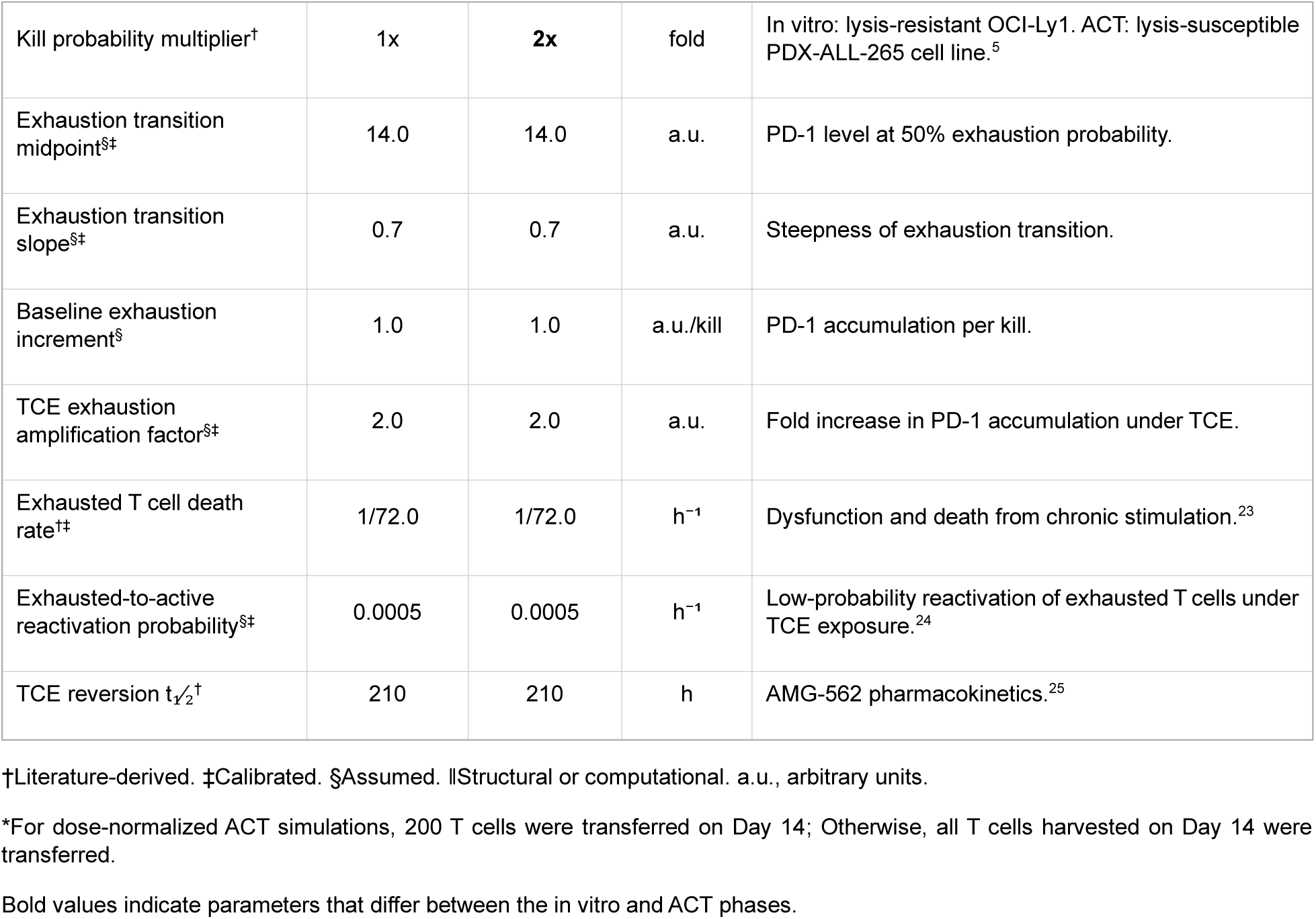
Model parameters, values, and sources.

Parameters marked ‡ in Table 1 were evaluated using a one-at-a-time (OAT) sensitivity analysis, in which each parameter was varied individually by ±100% of its calibrated or assumed value while all other parameters were held constant. The outcome evaluated was the percentage of exhausted CD8⁺ T cells under CONT and TFI schedules at Days 14, 21, and 28.

### Data Analysis, Visualization, and Reproducibility

To account for model stochasticity, each experimental condition was simulated across 50 independent iterations, each using a distinct predefined random seed and representing one independent stochastic realization of the model. Simulation outcomes are reported as mean ± standard deviation (SD), unless otherwise indicated. Statistical comparisons between groups were performed using the Mann-Whitney U test; P values were adjusted for multiple testing using the Bonferroni correction within each defined comparison family, as specified in the corresponding table or figure legend. Effect sizes were quantified using Cliff’s delta, and 95% confidence intervals for median differences were obtained via bootstrap resampling (10,000 iterations).

The model was implemented in Java (OpenJDK 21.0.2) using HAL,^19^ which requires JDK 1.8 or later. The full simulation code, parameter definitions, and run instructions are available at https://github.com/ninaobertopp/tce-tfi-abm-manuscript (MIT License, release v1.0.1). Claude (Anthropic) and ChatGPT (OpenAI) were used throughout manuscript preparation to assist with drafting and revising text, and drafting and debugging Java and Python code used in the model implementation and statistical analysis. The authors reviewed and edited all AI-assisted text and code. The authors take full responsibility for the accuracy, integrity, and conclusions of the manuscript.

## Results

### ABM Calibration Captures In Vitro Exhaustion Dynamics Under Continuous and Intermittent TCE Exposure

Previously, an in vitro lymphoma co-culture model of TCE therapy compared continuous 28-day exposure to the CD19×CD3 TCE AMG 562 (CONT), reflecting current clinical practice, with an intermittent regimen incorporating 7-day TFI.^5^ The study showed that T cells cultured under the TFI regimen exhibited greater proliferation and cytotoxicity, together with reduced exhaustion, compared with those cultured under CONT. We implemented the same CONT and TFI schedules and calibrated the ABM to the published percentages of exhausted CD8⁺ T cells at the corresponding time points (Fig. 2A–C).

**Figure 2.**
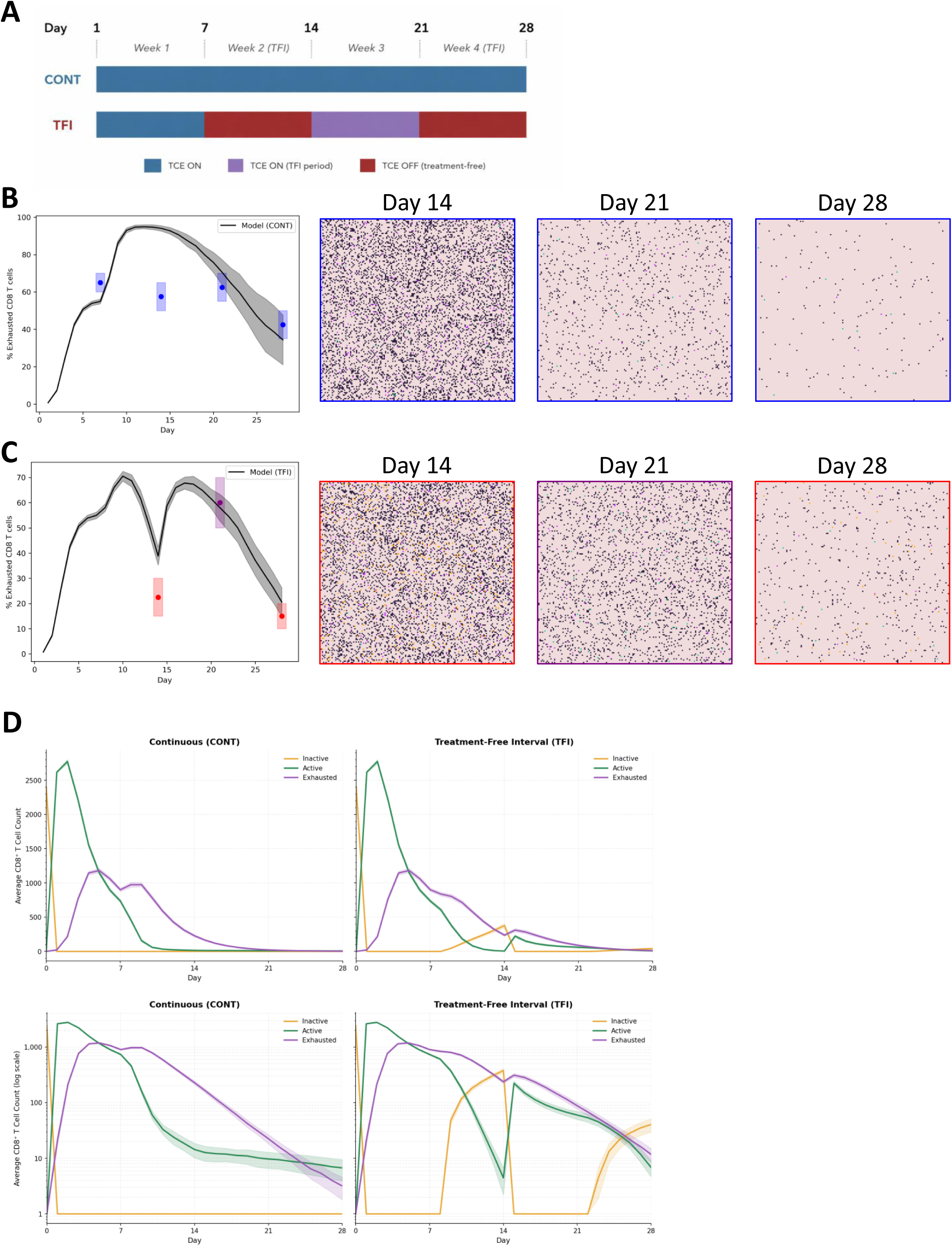
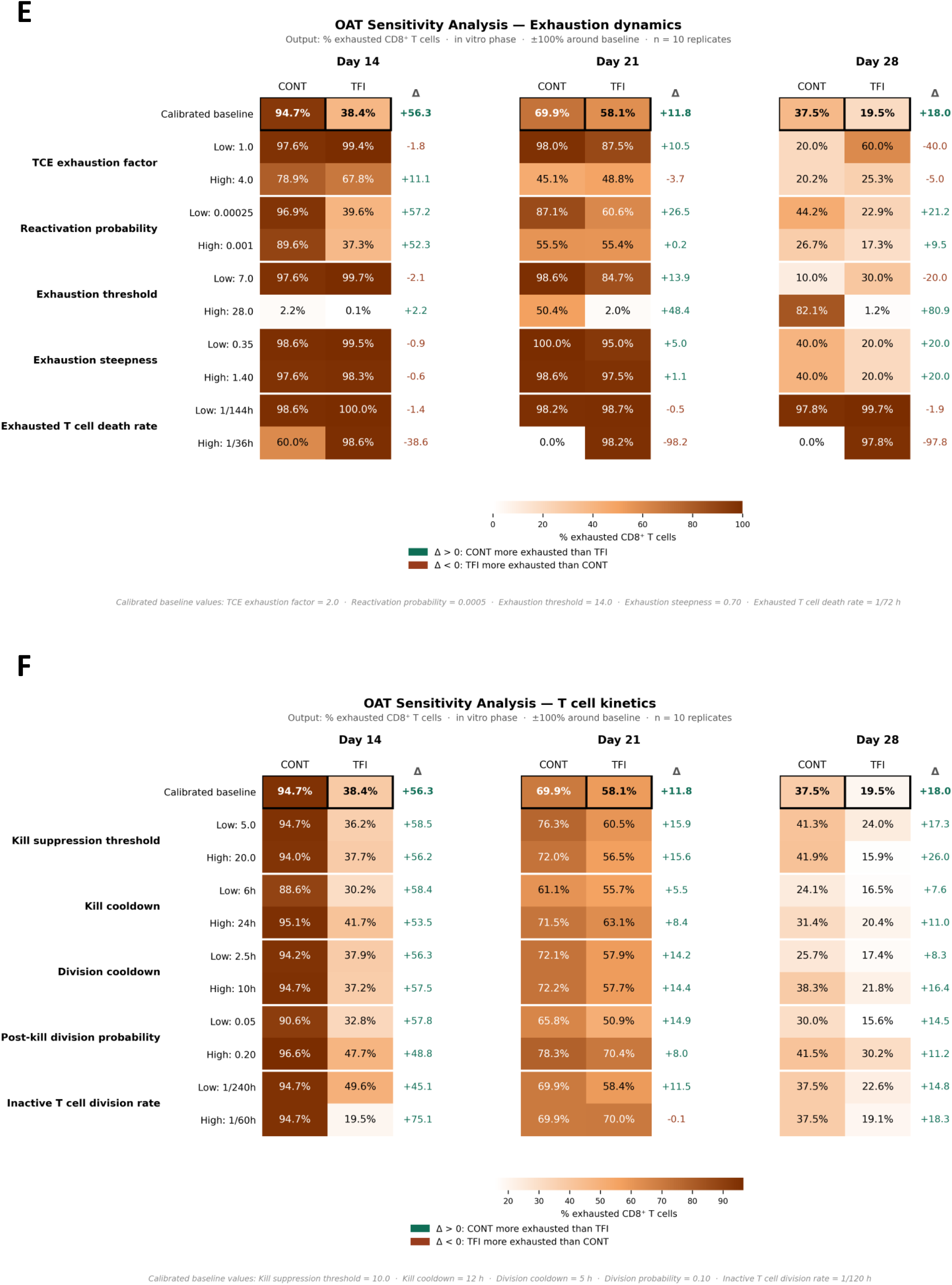
Agent-Based Model (ABM) Reproduces In Vitro Benefits of Intermittent T Cell Engager (TCE) Therapy. **(A)** Schematic of the two simulated TCE treatment schedules over the 28-day in vitro calibration phase. Top: continuous TCE administration (CONT), with treatment present from Day 1 through Day 28 (blue). Bottom: treatment-free interval (TFI) schedule, with TCE administered during Week 1 and Week 3 (blue/purple) and withdrawn during Week 2 and Week 4 (red), corresponding to the dosing structure reported in Philipp et al. **(B–C)** The black line shows the mean percentage of exhausted CD8⁺ T cells across replicate simulations, with gray shading indicating ±1 standard deviation (SD). Colored points indicate the median of the experimental measurements reported by Philipp et al. at matched timepoints, and shaded boxes denote the associated reported range. Blue points and boxes correspond to measurements under CONT; red points and boxes correspond to measurements following a TFI; purple points and boxes correspond to measurements obtained upon TCE re-exposure within the TFI schedule (Day 21). Representative spatial snapshots at Days 14, 21, and 28 are shown to the right of each panel, with tumor cells in black and T cells colored by functional state (inactive, orange; active, green; exhausted, purple); panel outline colors (blue, purple, red) indicate the corresponding treatment phase. **(D)** Simulated absolute CD8⁺ T cell counts by functional state (inactive, active, exhausted) over 28 days under CONT (left) and TFI (right) schedules, shown on linear (top) and logarithmic (bottom) scales. Lines represent the mean across replicates; shading denotes variability across replicates. **(E–F)** One-at-a-time (OAT) sensitivity analysis of exhaustion-related **(E)** and T cell kinetic **(F)** parameters. Heatmaps show the percentage of exhausted CD8⁺ T cells at Days 14, 21, and 28, averaged across n = 10 replicate simulations, for the calibrated baseline and after varying each parameter individually by ±100%. Green values indicate greater exhaustion under CONT than TFI, consistent with the experimental pattern reported by Philipp et al., whereas red values indicate the opposite trend.

Under both schedules, simulated exhaustion rose rapidly during the initial treatment period. Under CONT, high exhaustion was sustained before declining as exhausted cells were lost from the population. Under the TFI schedule, exhaustion fell substantially below CONT levels following TCE withdrawal, rose again upon re-exposure, and ultimately remained below CONT levels at Day 28 (Fig. 2B–C). Calibration showed closer quantitative agreement for the TFI schedule (RMSE = 5.10, n = 3 time points) than for CONT (RMSE = 14.74, n = 4 time points), although both reproduced the qualitative trends. Examination of the underlying T cell state composition showed that the TFI schedule preserved a larger non-exhausted T cell population that could respond to subsequent TCE re-exposure, whereas continuous exposure maintained a predominantly exhausted population (Fig. 2D).

To evaluate whether our findings were robust across parameter values, we varied each exhaustion-related and T cell kinetic parameter individually in an OAT sensitivity analysis (Fig. 2E–F). Across most parameter variations, exhaustion remained lower under TFI than CONT. The relationship between schedule and exhaustion was most sensitive to the exhausted T cell death rate, particularly when this rate was increased. Additional reversals at individual time points occurred when the TCE exhaustion factor was altered, the exhaustion threshold was lowered, or the exhaustion steepness was changed. Varying the reactivation probability did not reverse the relationship.

Taken together, the CD8⁺ T cell exhaustion dynamics simulated by our ABM were consistent with the calibration data from Philipp et al. We next examined how the resulting schedule-dependent T cell states affected antitumor activity following ACT.

### ABM Calibration Reproduces Anti-Tumor Benefits of Intermittent TCE Therapy

To determine whether the differences in T cell exhaustion observed during in vitro conditioning translated into differences in subsequent tumor control, we calibrated the ACT phase to the published in vivo experiments of Philipp et al. In that study, CONT- and TFI-conditioned T cells were harvested on Day 14 and transferred into tumor-bearing mice, which subsequently received AMG 562 on Days 2 and 9.^5^ To mirror this workflow, all simulated T cells present at the end of the 14-day in vitro phase were transferred to the ACT phase while retaining their individual phenotypes (Fig. 3A).

**Figure 3.**
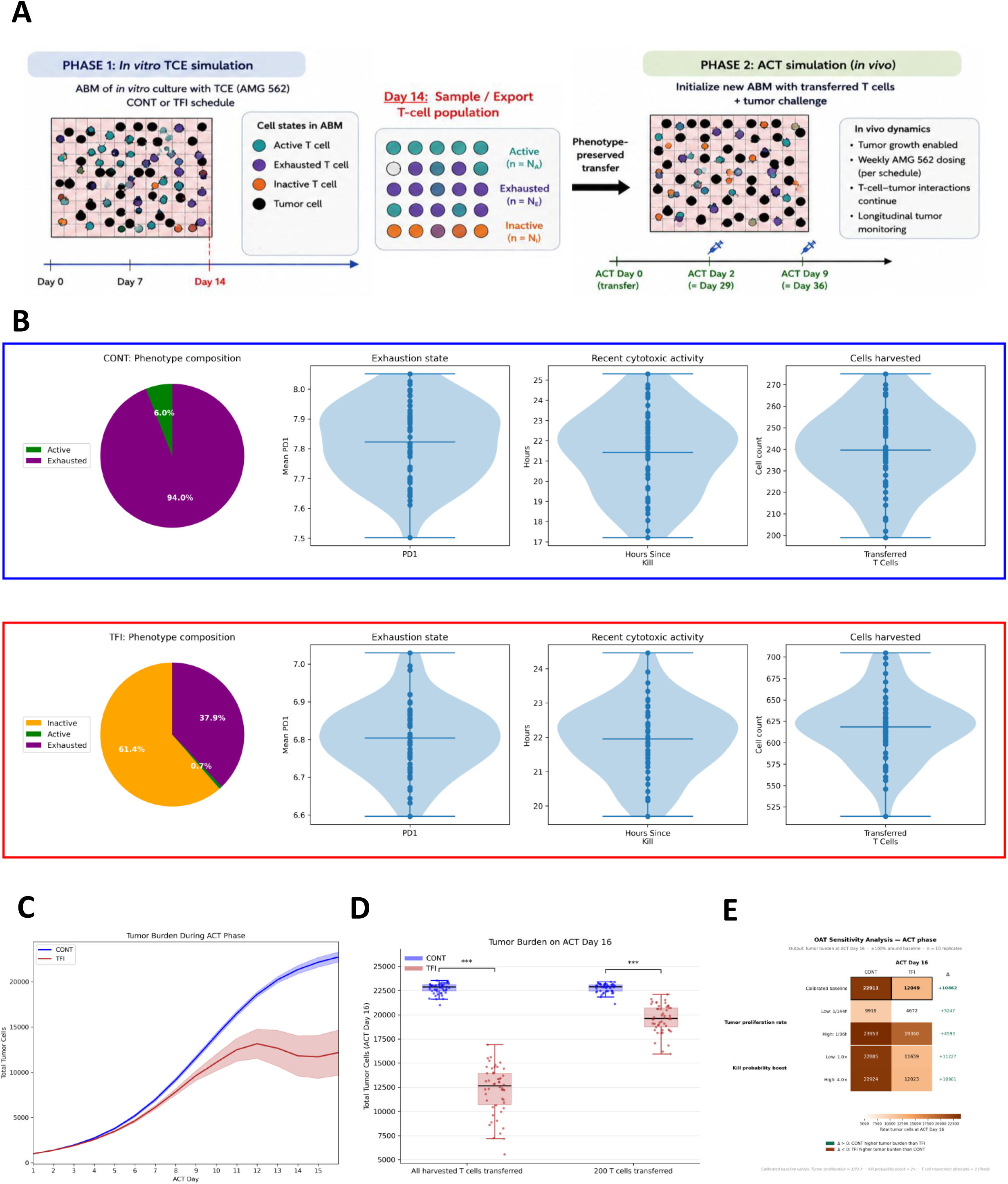
ABM Recapitulates Improved Tumor Control following Adoptive Cell Transfer (ACT) (A) Schematic of the two-phase simulation workflow. Phase 1: in vitro TCE conditioning under continuous exposure (CONT) or a treatment-free interval (TFI) schedule through Day 14. On Day 14, T cells are sampled for transfer while retaining their functional states (active, exhausted, or inactive). Phase 2: ACT with tumor growth and TCE administration on Days 2 and 9. (B) Composition and functional state of transferred T cell populations following CONT (top) or TFI (bottom) conditioning. Pie charts show the proportions of active, exhausted, and inactive T cells at transfer. Violin plots summarize PD-1 expression, hours since the most recent tumor-cell killing, and the number of transferred T cells across n = 50 replicate simulations. (C) Tumor burden during the ACT phase following transfer of CONT- or TFI-conditioned T cells, shown as total tumor cell counts over time (mean ± standard deviation [SD]). (D) Tumor burden on ACT Day 16 following transfer of all harvested T cells (left, as in C) or 200 T cells per replicate (right; S1 Fig.). Boxes show median and interquartile range; points represent individual replicates (n = 50). Mann-Whitney U test with Bonferroni correction. ***p < 0.001. (E) OAT sensitivity analysis of tumor burden on ACT Day 16, evaluating the effects of varying the tumor proliferation rate and T cell kill probability boost on total tumor cell counts under CONT and TFI conditions. The Δ column indicates the difference in tumor burden between CONT and TFI (CONT − TFI), with positive values (green) indicating greater tumor burden under CONT and negative values (red) indicating greater tumor burden under TFI.

At transfer, the CONT- and TFI-conditioned populations differed in phenotypic composition (Fig. 3B). The CONT-conditioned population was predominantly exhausted, whereas the TFI-conditioned population contained a larger inactive fraction and a smaller exhausted fraction. The mean exhaustion score was also significantly lower under TFI than CONT conditioning (6.80 ± 0.09 vs. 7.82 ± 0.13; Mann-Whitney U, p < 0.0001). In contrast, hours since last tumor-cell killing did not differ significantly between conditions (21.95 ± 1.03 vs. 21.42 ± 2.01 hours; p = 0.23).

TFI conditioning yielded more T cells by Day 14 than CONT conditioning (CONT: 239.64 ± 18.02 cells; TFI: 618.36 ± 38.69 cells). Consequently, approximately 2.6-fold more T cells were transferred from the TFI-conditioned population in the ACT simulation. Tumor burden increased more slowly following transfer of TFI-conditioned cells and remained lower than following transfer of CONT-conditioned cells (Fig. 3C), consistent with the experimental findings of Philipp et al.^5^

To determine whether the difference in transferred T cell number accounted for the improved tumor control, we repeated the ACT simulation with equal numbers of transferred T cells, randomly sampling 200 cells from each harvested population (S1 Fig. A, B). Even in the dose-normalized ACT simulation, tumor burden remained significantly lower following transfer of TFI-conditioned T cells (Day 16: p_adj = 2.57 × 10⁻¹⁷, Cliff’s δ = 0.99; Fig. 3D). Thus, the improved tumor control conferred by TFI conditioning did not depend on the greater number of cells transferred. At an equal transfer dose, the lower exhaustion score and larger inactive fraction of TFI-conditioned cells at harvest were sufficient to improve tumor control.

To evaluate whether our results were robust across ACT phase–specific parameter values, we performed an OAT sensitivity analysis by independently varying the ACT phase–specific tumor proliferation rate and kill-probability boost around their calibrated baseline values (Fig. 3E). Across all tested parameter variations, simulated tumor burden at ACT phase Day 16 remained lower following transfer of TFI-conditioned cells, demonstrating that this result was robust to the tested variation in tumor growth and cytotoxic activity.

### 7-Day TFI Outperforms Shorter and Continuous Schedules

After calibrating the ABM to the published experimental findings, we applied the ABM to examine how alternative TFI conditioning schedules affected ACT product quality and post-transfer tumor control. We compared six TCE conditioning schedules: continuous exposure (CONT), a Monday–Friday schedule (MO_FR), and four schedules with progressively shorter TFIs (77, TFI_5, TFI_3, and TFI_2; Fig. 4A). Across the 14-day conditioning phase, total TCE exposure ranged from 100% under CONT to 50% under TFI_7 (Fig. 4B).

**Figure 4.**
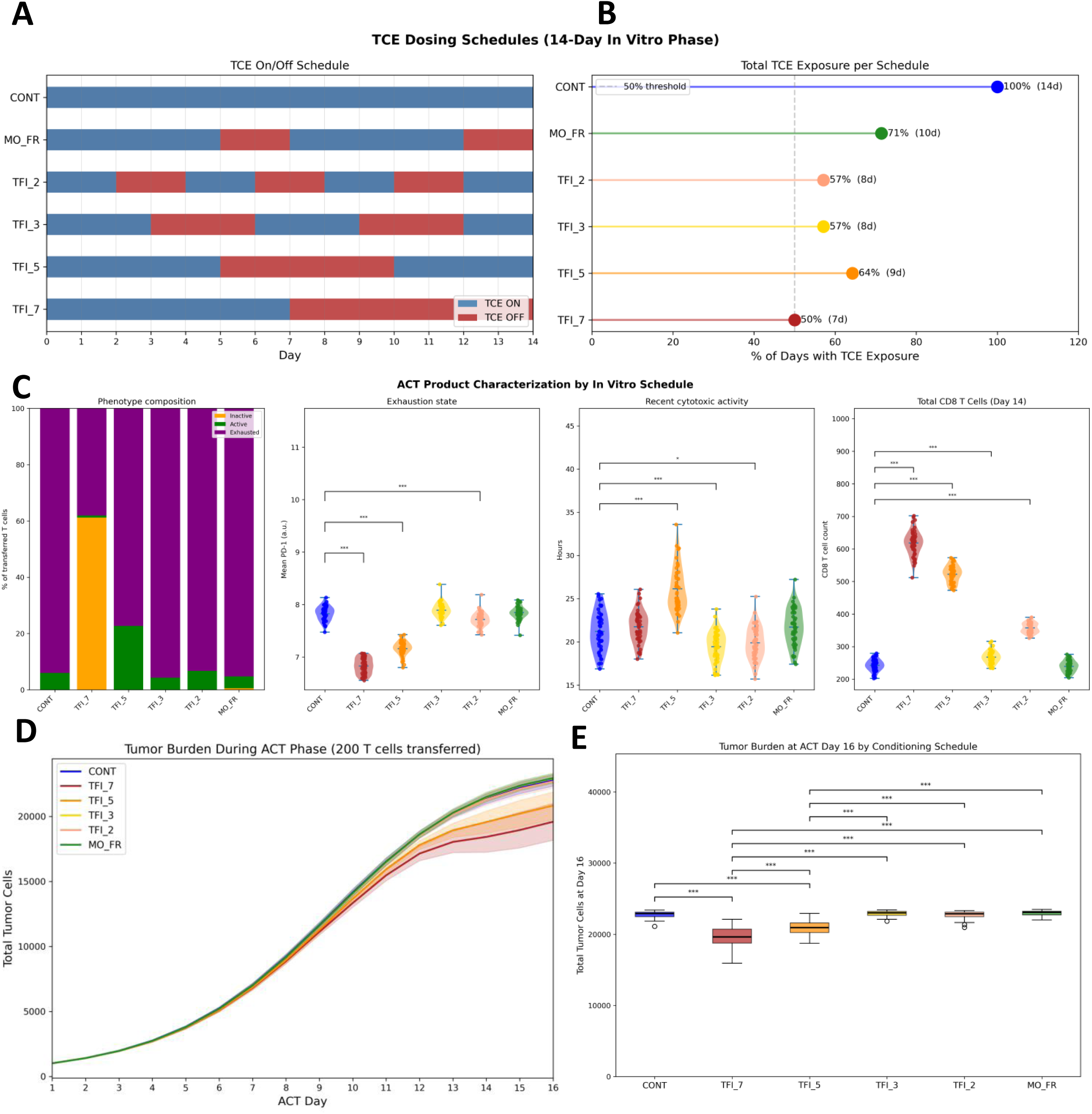
TCE Conditioning Schedules Shape ACT Product Phenotype and Post-Transfer Tumor Control. (A) TCE exposure schedules over the 14-day in vitro conditioning phase: continuous exposure (CONT), Monday–Friday exposure (MO_FR), and schedules with longest uninterrupted TFIs of 2, 3, 5, or 7 days (TFI_2, TFI_3, TFI_5, and TFI_7, respectively). Blue segments indicate days with TCE exposure; red segments indicate treatment-free days. (B) Total TCE exposure for each schedule, expressed as the percentage of the 14-day conditioning phase during which TCE was present. (C) Characterization of the resulting ACT products on Day 14, shown from left to right: phenotype composition (proportions of inactive, active, and exhausted CD8⁺ T cells), mean PD-1 expression, hours since the most recent tumor-cell kill, and total CD8⁺ T cell count. Comparisons with CONT were performed using the Mann–Whitney U test with Bonferroni correction within each metric; effect sizes (Cliff’s δ) and 95% confidence intervals are reported in S2 Table. (D) Simulated tumor burden over the 16-day ACT phase for each conditioning schedule following transfer of 200 T cells per condition. Lines represent mean tumor burden; shaded bands represent ±1 SD. (E) Tumor burden on ACT Day 16 across conditioning schedules. All 15 pairwise comparisons were performed using the Mann–Whitney U test with Bonferroni correction; effect sizes (Cliff’s δ) and 95% confidence intervals are reported in S3 Table. *p* < 0.05, \*\**p* < 0.001.

The different conditioning schedules generated distinct ACT products at harvest (Fig. 4C; S2 Fig. A). CONT, MO_FR, TFI_2, and TFI_3 all produced highly exhausted ACT products (>93% exhausted) with similarly elevated mean exhaustion scores. In contrast, TFI_5 reduced the exhausted fraction to 77.31 ± 3.32%, whereas TFI_7 produced the least exhausted ACT product, with only 38.04 ± 3.92% exhausted T cells and the lowest mean exhaustion score (6.83 ± 0.15). Notably, TFI_5 generated a substantially less exhausted ACT product than TFI_2 and TFI_3 despite including more cumulative TCE exposure (9 versus 8 days). This suggests that the temporal organization of TCE exposure, rather than cumulative exposure alone, influences T cell exhaustion at harvest.

The schedules also produced different total CD8⁺ T cell counts at Day 14. TFI_7, TFI_5, and TFI_2 each yielded significantly more cells than CONT (Cliff’s δ = −1.0 and p_adj < 5 × 10⁻¹⁷ for all three comparisons), whereas TFI_3 produced a smaller but significant increase (Cliff’s δ = −0.66, p_adj < 0.001). The CD8⁺ T cell count under MO_FR did not differ significantly from CONT (p_adj = 1.0). To separate the effects of ACT product quality from those of transfer dose, these ACT simulations used a normalized transfer dose of 200 T cells.

Differences in ACT product phenotype at harvest were reflected in tumor control following ACT (Fig. 4D–E; S2 Fig. B; S3 Table). TFI_7 significantly reduced tumor burden compared with all other schedules. Relative to CONT, TFI_7 produced a large reduction in tumor burden (Mann-Whitney U, p_adj < 0.001, Cliff’s δ = 0.99, 95% [2691, 3787] tumor cells). TFI_5 was the only other schedule that significantly reduced tumor burden relative to CONT (p_adj < 0.001, Cliff’s δ = 0.94), whereas MO_FR, TFI_2, and TFI_3 did not differ significantly from CONT, consistent with the high levels of T cell exhaustion observed after conditioning.

When T cell number was not normalized and all T cells present at the end of the in vitro phase were transferred, the advantage of TFI_7 became even more pronounced, while the overall ranking of the conditioning schedules was preserved (S3 Fig.). Together, these findings indicate that the benefit of TFI_7 was associated with reduced ACT product exhaustion and could not be explained by cumulative TCE exposure alone. Its advantage persisted after normalization of transfer dose and was further enhanced by its greater T cell yield.

### Placement of a 7-Day TFI Alters T Cell Phenotype Without Affecting Post-Transfer Tumor Control

Having found that, in the model, longer TFIs were associated with reduced T cell exhaustion and lower simulated tumor burden, and that this relationship could not be explained by cumulative TCE exposure alone, we next asked whether this benefit depends on the TFI’s placement within the conditioning schedule, or whether it holds regardless of when the rest period occurs. We therefore compared TFI_7 with TFI_7_offFirst, in which the same 7-day TFI was shifted to precede rather than follow TCE exposure while maintaining identical total TCE exposure (Fig. 5A–B).

**Figure 5.**
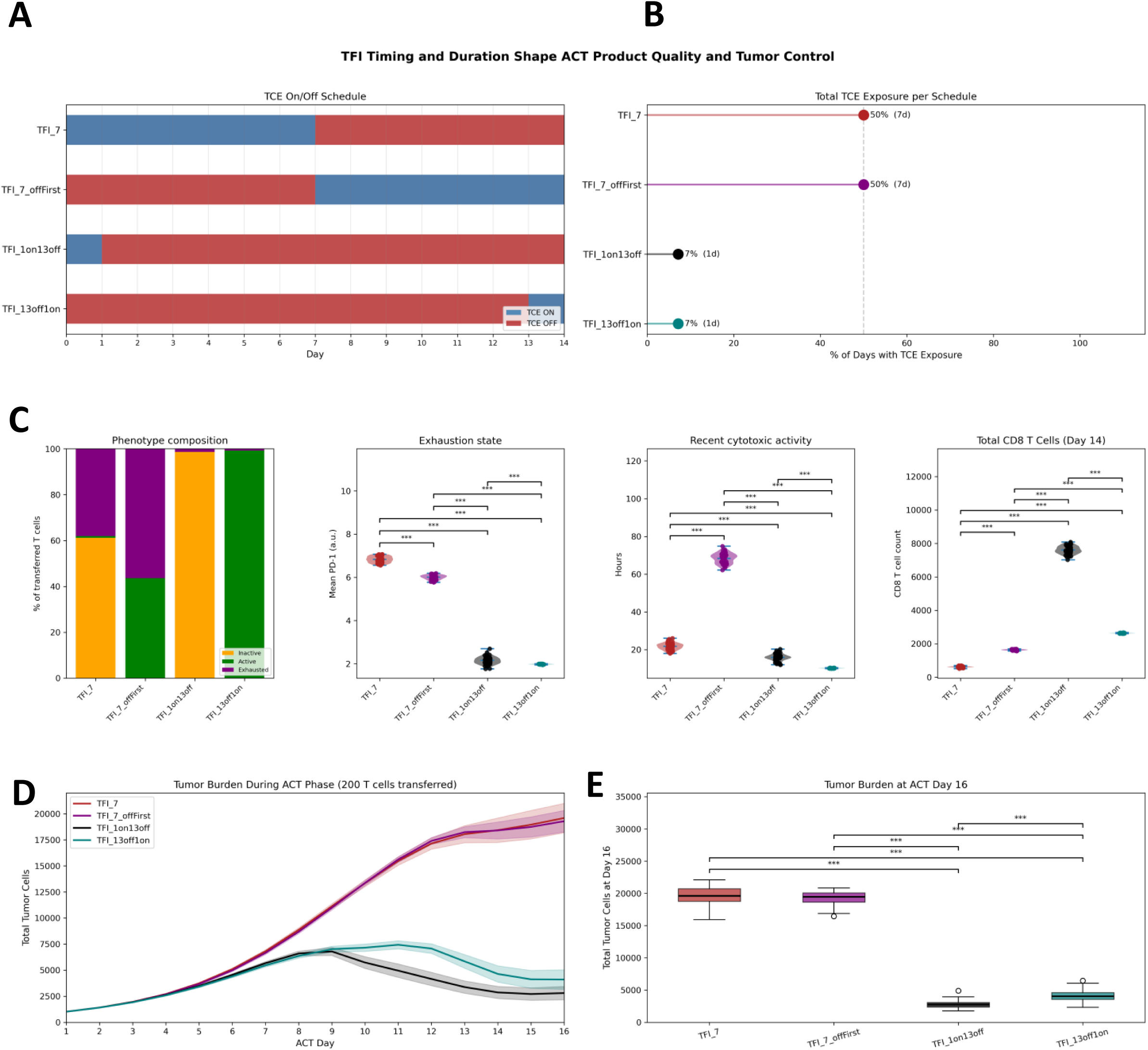
Effects of TFI Timing and Minimal TCE Exposure on ACT Product Phenotype and Post-Transfer Tumor Control. **(A)** TCE exposure schedules over the 14-day in vitro conditioning phase: TFI_7, with TCE exposure followed by a 7-day TFI; TFI_7_offFirst, with the 7-day TFI before TCE exposure; TFI_1on13off, with TCE exposure on the first day only; and TFI_13off1on, with TCE exposure on the final day only. Blue segments indicate days with TCE exposure; red segments indicate treatment-free days. **(B)** Total TCE exposure for each schedule, expressed as the percentage of the 14-day conditioning phase during which TCE was present. **(C)** Characterization of the resulting ACT products on Day 14, shown from left to right: phenotype composition (proportions of inactive, active, and exhausted CD8⁺ T cells), mean PD-1 expression, hours since the most recent tumor-cell kill, and total CD8⁺ T cell count. Pairwise comparisons were performed using the Mann–Whitney U test with Bonferroni correction within each metric; effect sizes (Cliff’s δ) and 95% confidence intervals are reported in S4 Table. **(D)** Simulated tumor burden over the 16-day ACT phase for each schedule following transfer of a fixed dose of 200 T cells per condition, sampled from the Day 14 population shown in (C). Lines represent mean tumor burden; shaded bands represent ±1 SD. **(E)** Tumor burden on ACT Day 16 across schedules. All six pairwise comparisons were performed using the Mann–Whitney U test with Bonferroni correction; effect sizes (Cliff’s δ) and 95% confidence intervals are reported in S5 Table. *p* < 0.05, \**p* < 0.01, \*\**p* < 0.001.

TFI_7 and TFI_7_offFirst produced significantly different T cell phenotypes at harvest despite identical total TCE exposure. TFI_7 yielded a predominantly inactive population (61.18 ± 3.88% inactive, 38.04 ± 3.88% exhausted, and 0.78 ± 0.59% active), with higher PD-1 expression (median, 6.83 vs. 6.01; Cliff’s δ = 1.0, p_adj = 4.21 × 10⁻¹⁷) and a more recent history of cytotoxic activity (median, 21.8 vs. 68.9 hours; Cliff’s δ = −1.0, p_adj = 4.24 × 10⁻¹⁷). By contrast, TFI_7_offFirst contained only active and exhausted T cells (43.62 ± 2.61% and 56.38 ± 2.61%, respectively), with no inactive cells (S4 Fig. A), and expanded to a larger total CD8⁺ T cell count by Day 14 (Cliff’s δ = −1.0, p_adj = 4.22 × 10⁻¹⁷; Fig. 5C; S4 Table). To isolate the effect of these phenotypic differences from that of T cell number, each ACT simulation was initialized with 200 transferred T cells.

Despite their phenotypic differences at harvest, tumor burden did not differ significantly between TFI_7 and TFI_7_offFirst on ACT Day 16 (Mann–Whitney U, p_adj = 0.87, Cliff’s δ = 0.17; S5 Table). This finding contrasts with the overall pattern across the schedules in Fig. 4, where a significantly higher exhausted fraction after conditioning was associated with significantly greater tumor burden following ACT (Spearman ρ = 0.93, p = 0.0067; S5 Fig.). However, the difference in exhausted fractions between TFI_7 and TFI_7_offFirst was substantially smaller than that between TFI_7 and TFI_5 (38.04 ± 3.88% vs. 56.38 ± 2.61% and 38.04 ± 3.88% vs. 77.31 ± 3.32%, respectively). Their T cell state compositions also became increasingly similar following TCE administration during the ACT phase (S4 Fig. B). Together, these findings suggest that, although the two 7-day schedules produced significantly different phenotypes at harvest, these differences were not sufficiently large or persistent to produce a detectable difference in post-transfer tumor control. Thus, phenotype at harvest alone may not fully predict subsequent antitumor activity.

### One-Day TCE Schedules Reveal a Limitation of TFI Testing In Vitro

Among the initially tested conditioning schedules, TFI_7 produced the least exhausted ACT product and the lowest post-transfer tumor burden. Because TFI_7 also had the lowest total TCE exposure, we next asked whether further reducing exposure would improve tumor control. We evaluated two schedules with only one day of TCE exposure during the 14-day conditioning period, administered either on the first day (TFI_1on13off) or immediately before harvest (TFI_13off1on). Both schedules substantially reduced simulated tumor burden compared with the 7-day TFI schedules (Mann–Whitney U test, p_adj < 5 × 10⁻¹⁷ and Cliff’s δ ≥ 0.98 for all comparisons; S5 Table), consistent with greater preservation of the T cell population during conditioning.

At harvest, TFI_1on13off produced an almost entirely inactive ACT product (98.6 ± 0.8% inactive and 1.4 ± 0.8% exhausted), whereas TFI_13off1on produced an almost exclusively active T cell product. TFI_1on13off also generated a substantially larger Day 14 CD8⁺ T cell population than every other schedule tested (7,610.0 ± 239.0 cells, compared with 618.0 ± 38.4 for TFI_7, 1,636.6 ± 31.3 for TFI_7_offFirst, and 2,636.0 ± 15.1 for TFI_13off1on; all Cliff’s δ ≥ 0.73 and p_adj < 5 × 10⁻¹⁷; Fig. 5C; S4 Table), consistent with minimal TCE-driven exhaustion and attrition across most of the conditioning period. However, the number of transferred T cells was fixed at 200 across ACT simulations, preventing the larger TFI_1on13off population from directly conferring a numerical advantage after transfer.

Unlike the comparison between the 7-day schedules, the timing of the single TCE exposure influenced simulated tumor control during the ACT phase. TFI_1on13off achieved significantly lower tumor burden than TFI_13off1on at Day 16 (p_adj = 3.6 × 10⁻¹⁰, Cliff’s δ = −0.76). This difference occurred despite significantly higher PD-1 expression under TFI_1on13off (p_adj = 1.85 × 10⁻⁹, Cliff’s δ = 0.73; S4 Table), suggesting that, within the model, the functional-state composition of the transferred T cell population may have been more closely related to subsequent tumor control than PD-1 expression alone.

Importantly, the apparent superiority of the one-day TCE schedules should not be interpreted as evidence that they are clinically optimal. Instead, these schedules expose a potential limitation of the preclinical approach used by Philipp et al.^5^ and reproduced by the ABM. In both the preclinical experiment and our computational model, TFIs are applied during in vitro conditioning in the absence of a growing tumor. This design allows prolonged treatment withdrawal to reduce T cell exhaustion without accounting for concurrent tumor progression. If applied directly in vivo, a 13-day treatment-free interval would likely permit tumor growth that could offset the apparent advantage observed here. This limitation reflects current constraints on preclinical models for directly evaluating TFI schedules in vivo.^33,34^

## Discussion

We developed an ABM of TCE therapy calibrated to key experimental observations reported by Philipp et al. The model reproduced their central finding that interrupting continuous TCE exposure with TFIs reduces T cell exhaustion and improves subsequent antitumor activity.^5^ TFI conditioning also increased simulated T cell yield. After equalizing the transferred cell dose, the improvement in tumor control persisted, showing that cell number alone did not explain the benefit. We then used the model to investigate how TFI duration and timing affect T cell phenotype and tumor control following ACT.

Extending beyond the experimentally tested 7-day TFI schedule, our simulations showed that both the duration and timing of TFIs affected the resulting T cell populations and subsequent tumor control. Among the schedules evaluated, TFI_7 generated the least exhausted T cell population and achieved the greatest tumor control following adoptive transfer, TFI_5 provided an intermediate benefit, and shorter interruptions produced outcomes similar to continuous exposure.

Moving the same 7-day TFI from the second half to the first half of the conditioning period (TFI_7_offFirst) produced markedly different T cell populations at harvest, yet post-transfer tumor control was essentially unchanged. Following transfer, both populations were exposed to the same AMG 562 schedule and converged in their functional-state dynamics over the course of the ACT phase. This suggests that phenotype at harvest alone may not fully predict post-transfer tumor control, because T cell functional states continue to change following TCE re-exposure. In these simulations, initially distinct populations became increasingly similar during the ACT phase and achieved comparable tumor control.

Extending the analysis to schedules with only one day of TCE exposure during the 14-day in vitro phase produced even less exhausted ACT products and greater post-transfer tumor control. However, the strong performance of these schedules also revealed an important limitation of the sequential conditioning and ACT design. During in vitro conditioning, tumor cells are experimentally replenished rather than continuously growing as they would in vivo. Consequently, prolonged TFIs can reduce chronic T cell stimulation without the trade-off of tumor progression that would occur during treatment withdrawal in vivo.

Several additional limitations should be considered when interpreting these findings. First, calibration produced a closer quantitative fit to the TFI data than to the CONT data. This difference reflects the difficulty of calibrating a stochastic, rule-based ABM, in which nonlinear interactions among model components complicate the relationship between parameter values and model outcomes.^35^

Second, direct evaluation of TCE schedules in immunocompetent mice remains challenging because these molecules are engineered to engage human, not murine, CD3, and human PDX tumors require immunodeficient hosts, preventing their evaluation within a syngeneic, fully immunocompetent tumor–immune system.^34,36^ Consistent with these experimental constraints, Philipp et al. used an experimental design in which healthy-donor T cells were first conditioned with AMG 562 in vitro and subsequently transferred into NSG mice bearing PDX-ALL.^5^ Their experiment tested the in vivo activity of differently conditioned T cells rather than comparing TFI schedules administered directly to tumor-bearing mice. The current ABM reproduces this experimental design and consequently inherits its limitations. The separation of in vitro conditioning from ACT does not reflect conventional TCE therapy, which engages endogenous T cells in vivo. Nevertheless, this two-phase structure resembles ACT modalities such as CAR T-cell therapy and tumor-infiltrating lymphocytes (TIL) therapy, in which T cells undergo ex vivo processing and expansion before adoptive transfer. With appropriate adaptations, the ABM presented here could provide a foundation for examining how T cell number and phenotype at transfer influence subsequent antitumor activity in these settings.

Third, the pharmacokinetic assumptions used to model AMG 562 introduce additional uncertainty. The currently FDA-approved TCE for B-ALL, blinatumomab, has a short half-life of approximately 2 hours and therefore requires continuous intravenous infusion.^37^ To overcome the technical challenges associated with maintaining continuous exposure during experimental TFI studies, Philipp et al. used AMG 562^5^, a half-life-extended CD19×CD3 bispecific molecule with a reported half-life of approximately 210 hours. To reflect its prolonged persistence in our model, we incorporated a probabilistic decay function based on the reported in vivo half-life of AMG 562.^25^ This allowed us to simulate a gradual decline in TCE activity and T cell reversion dynamics during TFIs. However, pharmacokinetic properties such as clearance and half-life do not translate directly between in vivo and in vitro systems because in vitro cultures lack the corresponding physiological elimination processes.^38^ As a result, the actual half-life of AMG 562 in the in vitro setting may be longer than modeled. Because the model’s TFI dynamics are calibrated to AMG 562’s substantially longer half-life, the simulated schedules and their effects on T cell reversion cannot be directly extrapolated to blinatumomab, whose rapid clearance would be expected to produce much faster T cell reversion following treatment withdrawal. Accordingly, clinical translation of the simulated schedules will require consideration of formulation-specific exposure kinetics, particularly given the ongoing development of subcutaneous TCE formulations.^39^

Combining immunotherapy with other treatment modalities has improved clinical outcomes across multiple cancer types.^40^ Building on our finding that treatment timing influences T cell state and antitumor activity, an expanded version of the ABM could be used to investigate how TCEs should be sequenced with other treatments. One relevant combination is TCE therapy with PD-1/PD-L1 blockade, for which the timing of checkpoint inhibition may alter the balance between T cell activation and exhaustion. This question may become increasingly important as multispecific TCEs with integrated checkpoint inhibition are developed.^41^ Radiation is another relevant combination for immunotherapy because it can enhance T cell-mediated antitumor responses.^42^ An expanded ABM could investigate the optimal sequencing of TCE and radiation therapy, particularly in solid tumors, where limited T cell trafficking, local immunosuppression, and tumor and antigen heterogeneity restrict TCE efficacy.^43^ By explicitly representing these spatial tumor–immune interactions, ABMs could help determine how different locally delivered radiation schedules alter responses to TCE therapy.^11^ Consistent with this approach, an ABM has already been used to examine how spatially heterogeneous radiation exposure affects tumor–immune dynamics and treatment response.^8^

Our ABM reproduced the central experimental finding of Philipp et al. that TFIs can preserve T cell function during otherwise continuous TCE exposure.^5^ We then extended the analysis beyond the experimentally evaluated schedules and found that both TFI duration and timing influenced T cell phenotype and subsequent tumor control. Longer uninterrupted TFIs were associated with less exhaustion and improved post-transfer tumor control. However, the apparent benefit of schedules containing only one day of TCE exposure highlighted a potential limitation of the sequential conditioning and ACT design, which cannot capture the cost of tumor progression during treatment withdrawal. These findings motivate the development of an in vivo ABM that models T cell exhaustion, TCE exposure, and tumor progression concurrently, thereby capturing the trade-off between preserving T cell function during treatment withdrawal and maintaining tumor control. Integrating such a model with experimental validation could help identify effective intermittent TCE regimens.

## Author Contributions

N.O.: Conceptualization, Methodology, Formal analysis, Investigation, Visualization, Writing – original draft.

M.F.: Methodology, Writing – review & editing.

D.B.: Supervision, Writing – review & editing.

S.P.-T.: Funding acquisition

## Data Availability

The full simulation code, parameter definitions, and run instructions are available at https://github.com/ninaobertopp/tce-tfi-abm-manuscript (MIT License, release v1.0.1), archived at Zenodo (DOI: 10.5281/zenodo.22212428).

