## Supplements for "Treatment-free interval duration during T-cell engager conditioning shapes simulated T-cell exhaustion and post-transfer tumor control"

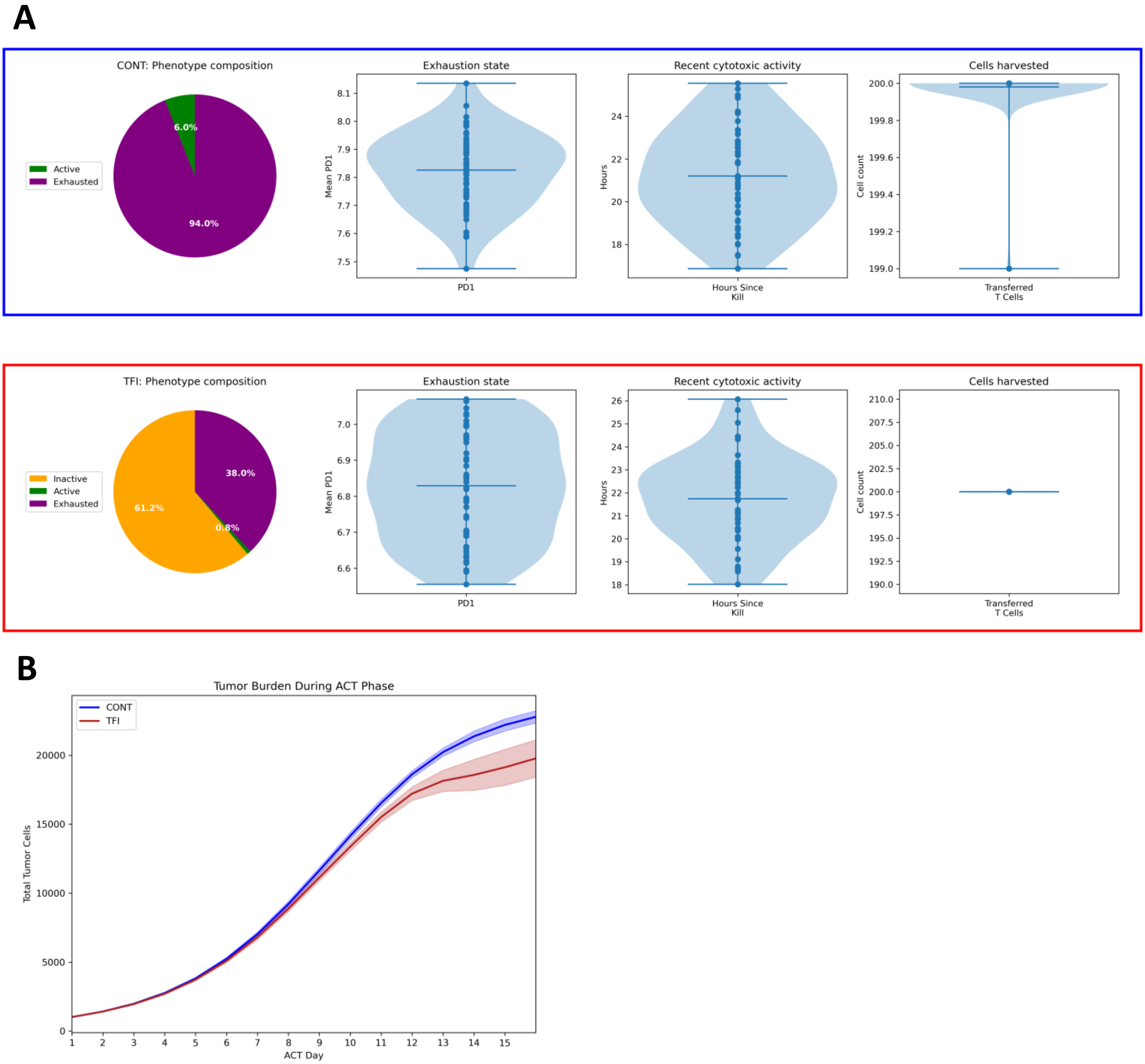
A
CONT: Phenotype composition
Active
Exhausted
6.0%
94.0%
Exhaustion state
Mean PD1
PD1
Recent cytotoxic activity
Hours
Hours Since Kill
Cells harvested
Cell count
Transferred T Cells
TFI: Phenotype composition
Inactive
Active
Exhausted
38.0%
61.2%
0.8%
Exhaustion state
Mean PD1
PD1
Recent cytotoxic activity
Hours
Hours Since Kill
Cells harvested
Cell count
Transferred T Cells
B
Tumor Burden During ACT Phase
CONT
TFI
Total Tumor Cells
ACT Day

**S1 Fig. ACT Product Phenotype and Post-Transfer Tumor Control Under CONT and TFI TCE Conditioning with ACT Fixed at 200 T Cells**

**(A)** Characterization of the ACT products on Day 14 following continuous TCE exposure (CONT; top) or a treatment-free interval schedule (TFI; bottom). Pie charts show the proportions of active, inactive, and exhausted $CD8^+$ T cells. Violin plots show mean PD-1 expression, hours since the most recent tumor-cell kill, and the total number of T cells harvested before dose normalization. For the ACT phase, transfer was limited to 200 T cells per replicate.

**(B)** Simulated tumor burden over the 16-day ACT phase following transfer of 200 CONT- or TFI-conditioned T cells. Lines represent mean tumor burden; shaded bands represent ±1 SD.

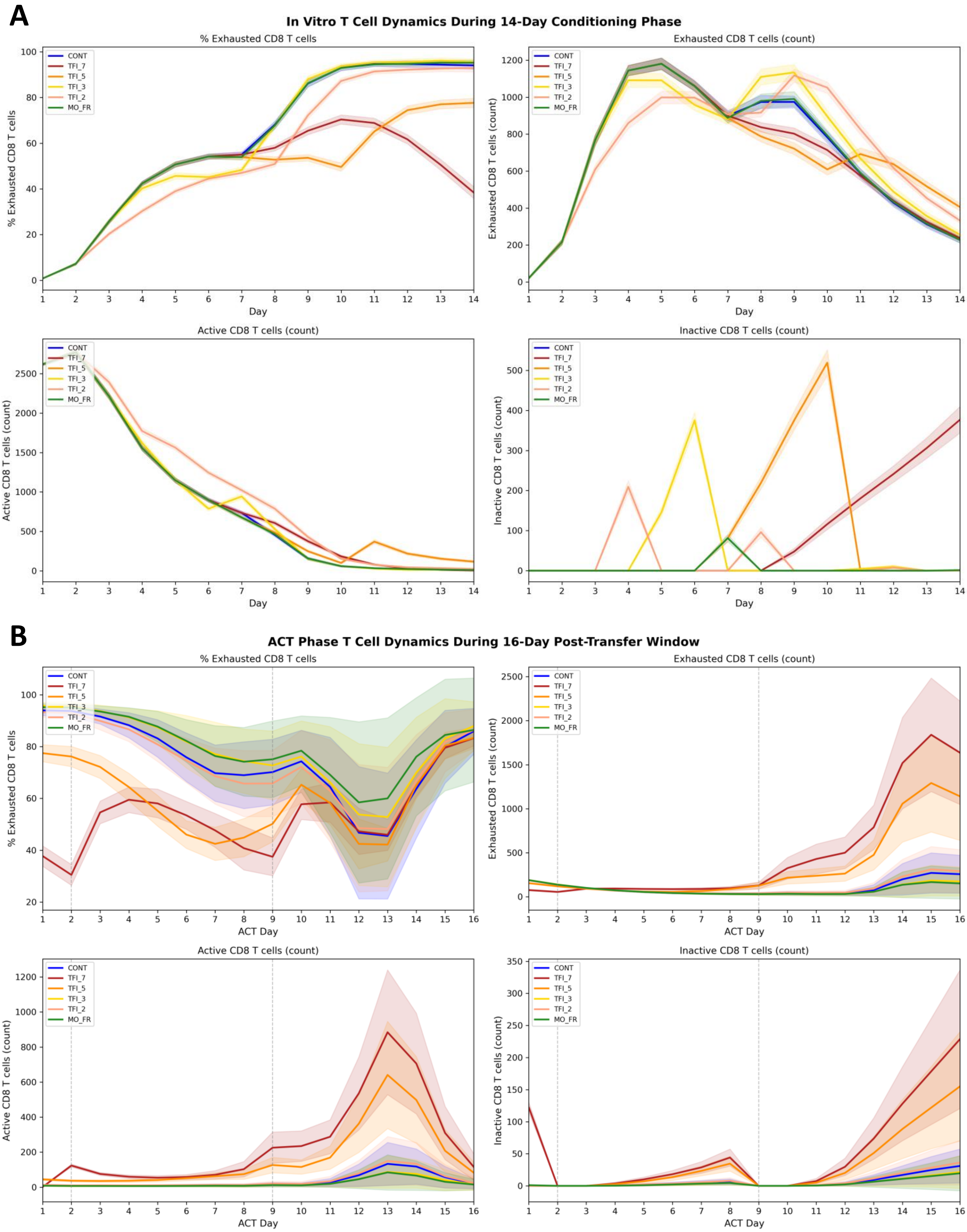
A
In Vitro T Cell Dynamics During 14-Day Conditioning Phase
% Exhausted CD8 T cells
Exhausted CD8 T cells (count)
Active CD8 T cells (count)
Inactive CD8 T cells (count)
Day
CONT
TFI_7
TFI_5
TFI_3
TFI_2
MO_FR
B
ACT Phase T Cell Dynamics During 16-Day Post-Transfer Window
% Exhausted CD8 T cells
Exhausted CD8 T cells (count)
Active CD8 T cells (count)
Inactive CD8 T cells (count)
ACT Day

**S2 Fig. T Cell Dynamics During Conditioning and ACT Across TCE Schedules Without Transfer-Dose Normalization**

**(A)** In vitro T cell dynamics over the 14-day conditioning phase for each TCE schedule (CONT, TFI_7, TFI_5, TFI_3, TFI_2, and MO_FR), shown clockwise from the top left: percentage of exhausted $CD8^+$ T cells, exhausted $CD8^+$ T cell count, inactive $CD8^+$ T cell count, and active $CD8^+$ T cell count. **(B)** T cell dynamics over the 16-day ACT phase following transfer of all T cells harvested at the end of conditioning, shown using the same four metrics. Lines represent mean values; shaded bands represent ±1 SD.

A

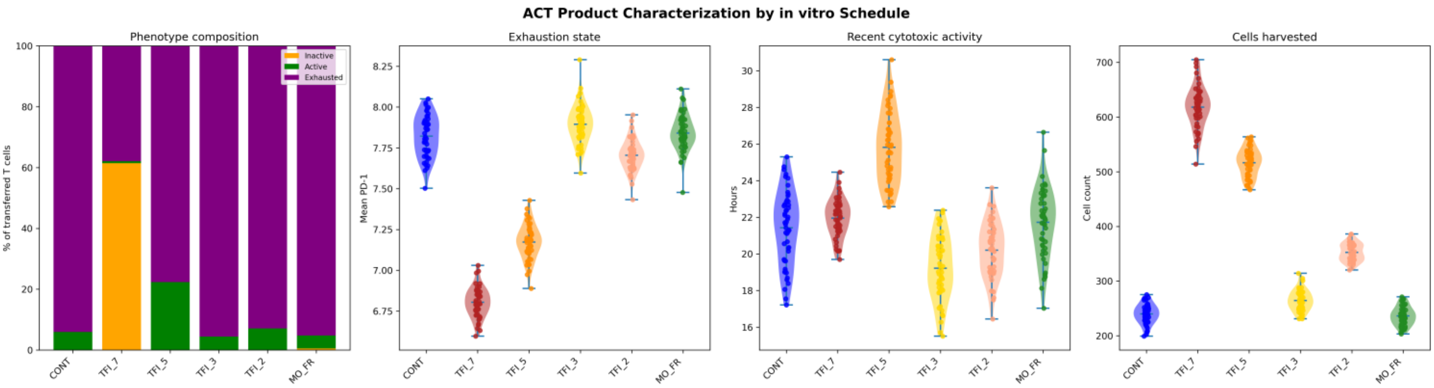


B

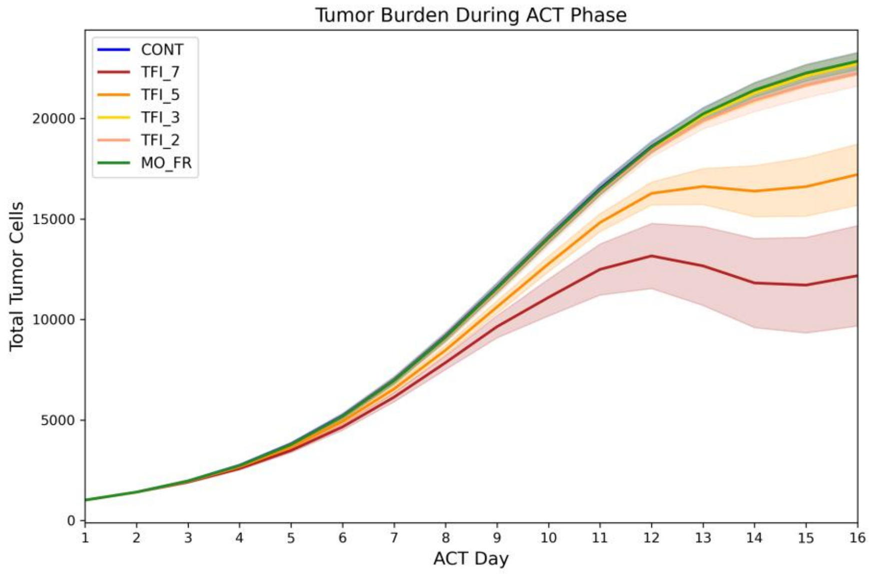


**S3 Fig. ACT Product Phenotype and Post-Transfer Tumor Burden Across Six Conditioning Schedules Without Transfer-Dose Normalization**

**(A)** Characterization of the ACT products on Day 14 across the six in vitro conditioning schedules (CONT, TFI_7, TFI_5, TFI_3, TFI_2, and MO_FR), shown from left to right: phenotype composition (proportions of inactive, active, and exhausted CD8$^{+}$ T cells), mean PD-1 expression, hours since the most recent tumor-cell kill, and total T cells harvested. **(B)** Simulated tumor burden over the 16-day ACT phase following transfer of all T cells harvested under each conditioning schedule. Lines represent mean tumor burden; shaded bands represent ±1 SD.

**A**

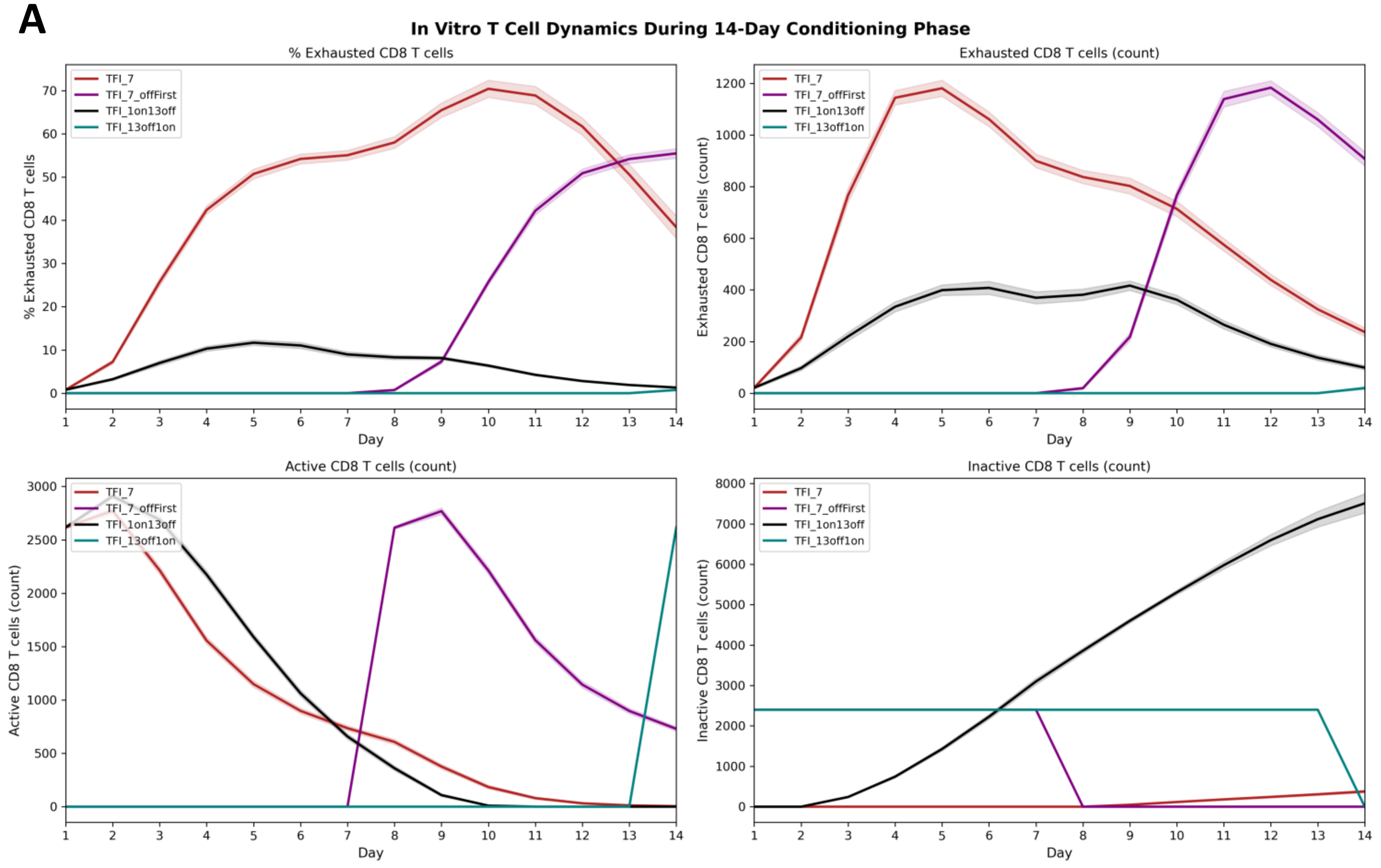
In Vitro T Cell Dynamics During 14-Day Conditioning Phase
% Exhausted CD8 T cells
Exhausted CD8 T cells (count)
Active CD8 T cells (count)
Inactive CD8 T cells (count)
Day
TFI_7
TFI_7_offFirst
TFI_1on13off
TFI_13off1on

**B**

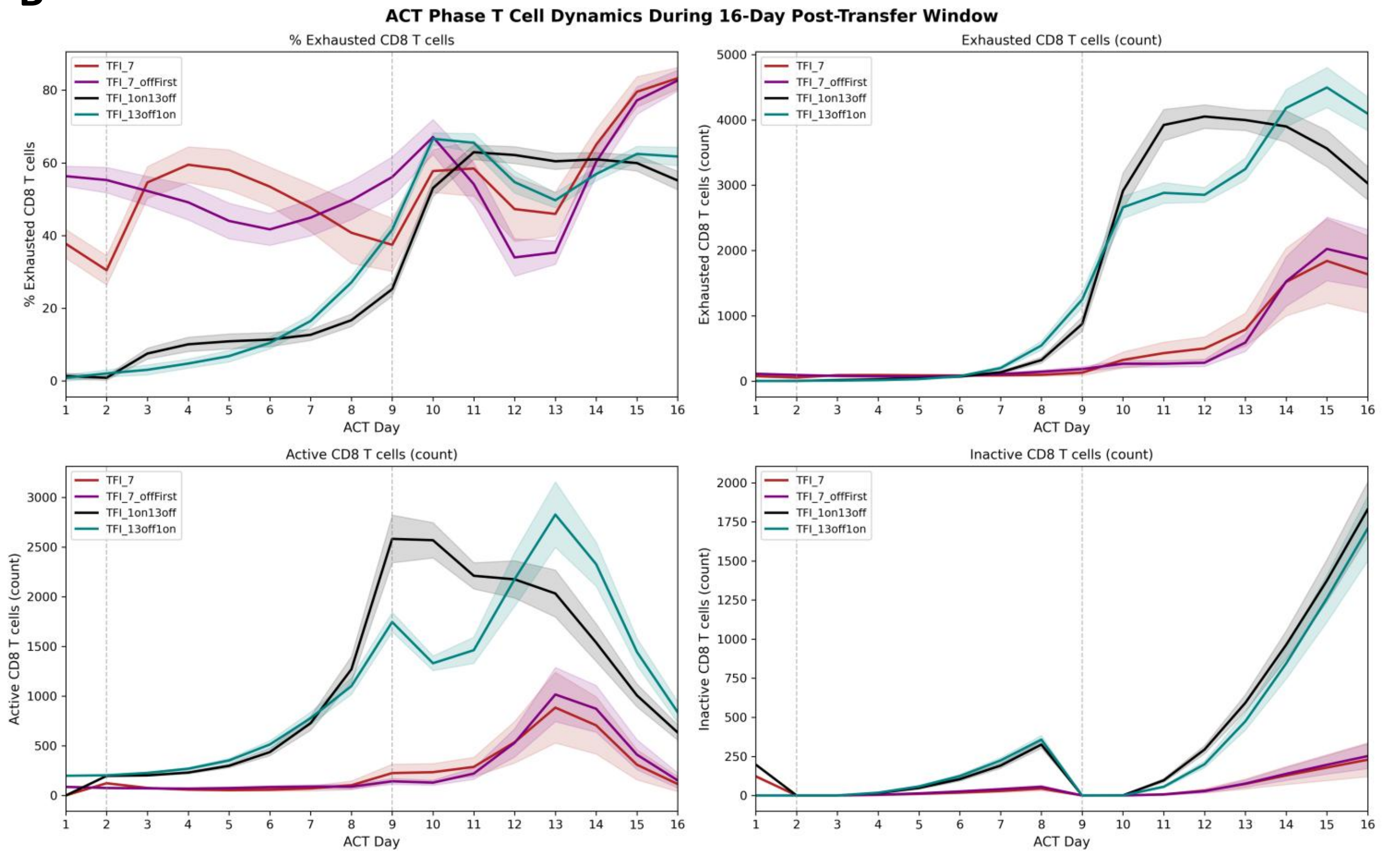
ACT Phase T Cell Dynamics During 16-Day Post-Transfer Window
% Exhausted CD8 T cells
Exhausted CD8 T cells (count)
Active CD8 T cells (count)
Inactive CD8 T cells (count)
ACT Day
TFI_7
TFI_7_offFirst
TFI_1on13off
TFI_13off1on

**S4 Fig. T Cell Dynamics Across TFI Timing and Minimal TCE Exposure Schedules**

**(A)** In vitro T cell dynamics over the 14-day conditioning phase for TFI_7, TFI_7_offFirst, TFI_1on13off, and TFI_13off1on, shown clockwise from the top left: percentage of exhausted $CD8^+$ T cells, exhausted $CD8^+$ T cell count, inactive $CD8^+$ T cell count, and active $CD8^+$ T cell count. **(B)** T cell dynamics over the 16-day ACT phase following transfer of a fixed dose of 200 T cells per schedule, shown using the same four metrics. Lines represent mean values; shaded bands represent ±1 SD.

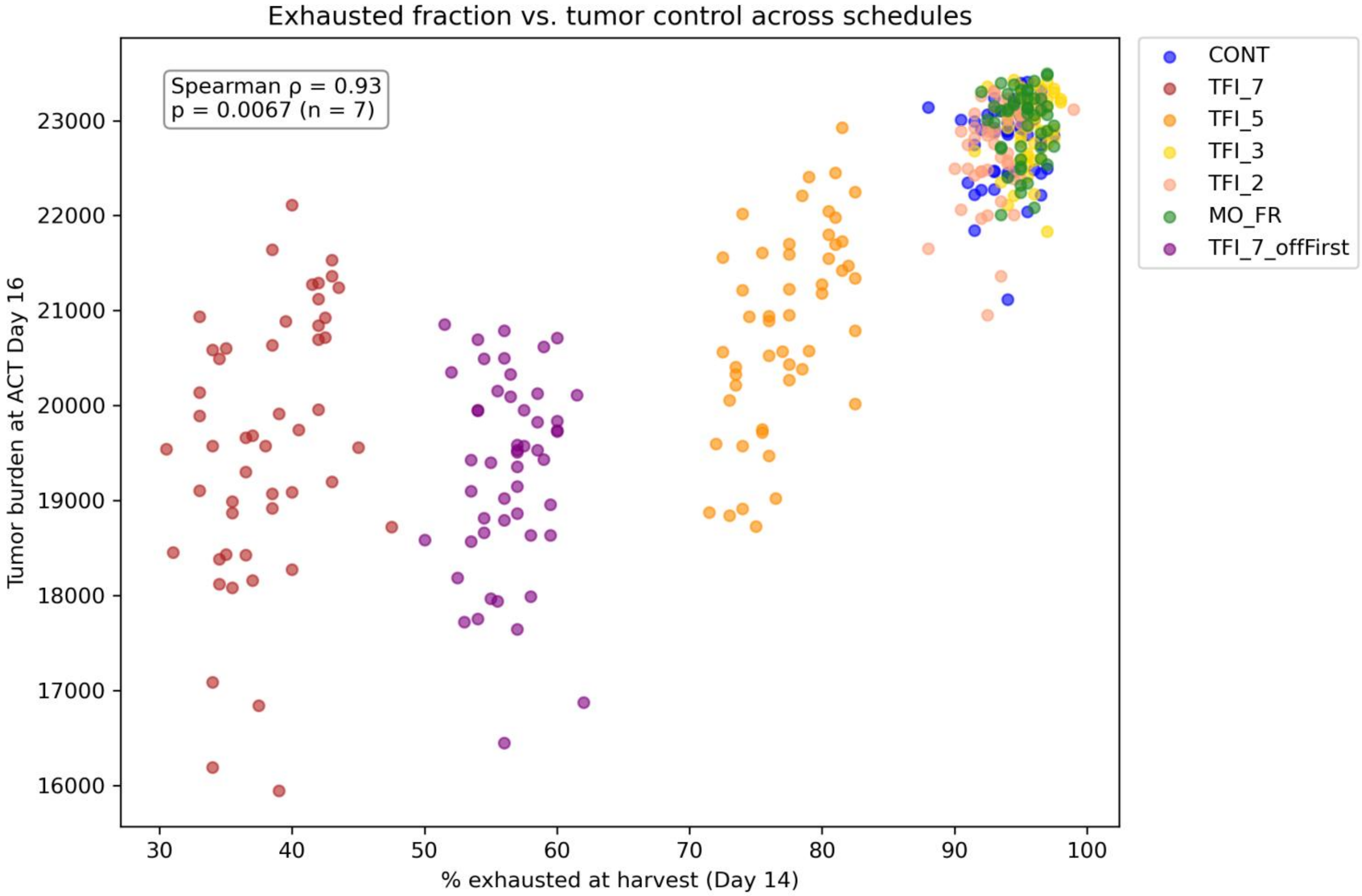


**S5 Fig. Exhausted Fraction at Harvest Correlates with Post-Transfer Tumor Burden Across Conditioning Schedules**

The proportion of exhausted $CD8^+$ T cells in the ACT product on Day 14 (x-axis) is plotted against simulated tumor burden on ACT Day 16 (y-axis) for each replicate across seven conditioning schedules: the six schedules evaluated in Fig. 4 (CONT, MO_FR, TFI_2, TFI_3, TFI_5, and TFI_7) and TFI_7_offFirst. Each point represents one simulation replicate; colors indicate the conditioning schedule. The Spearman correlation was calculated using the mean values for each schedule rather than individual replicate values (n = 7; ρ = 0.93, *p* = 0.0067; exact permutation test).

**S1 Table.** Schedule Design Parameters, ACT Product Phenotype, and Day 16 Tumor Burden Across TCE Conditioning Schedules

| Schedule | Total exposure (%) | Longest off-interval (days) | TCE on before harvest (days) | TCE off before harvest (days) | Inactive (%) | Active (%) | Exhausted (%) | Mean PD-1 | Hours since last kill | Day 16 tumor burden vs. CONT (Mann-Whitney U, Bonferroni-adjusted) |
|---|---|---|---|---|---|---|---|---|---|---|
| **CONT** | 100 | 0 | 14 | 0 | 0.00 ± 0.00 | 6.01 ± 1.90 | 93.99 ± 1.90 | 7.83 ± 0.14 | 21.20 ± 2.23 | Reference |
| **MO_FR** | 71 | 2 | 0 | 2 | 0.53 ± 0.48 | 4.26 ± 1.26 | 95.21 ± 1.36 | 7.85 ± 0.13 | 21.70 ± 2.15 | Not significant (p_adj = 0.41) |
| **TFI_2** | 57 | 2 | 2 | 0 | 0.00 ± 0.00 | 6.73 ± 1.93 | 93.27 ± 1.93 | 7.72 ± 0.13 | 19.88 ± 1.90 | Not significant (p_adj = 1.00) |
| **TFI_3** | 57 | 3 | 2 | 0 | 0.00 ± 0.00 | 4.27 ± 1.35 | 95.73 ± 1.35 | 7.89 ± 0.13 | 19.45 ± 1.78 | Not significant (p_adj = 1.00) |
| **TFI_5** | 64 | 5 | 4 | 0 | 0.00 ± 0.00 | 22.69 ± 3.32 | 77.31 ± 3.32 | 7.16 ± 0.13 | 26.13 ± 2.63 | Significantly lower (p_adj = $8.96 \times 10^{-15}$) |
| **TFI_7** | 50 | 7 | 0 | 7 | 61.18 ± 3.92 | 0.78 ± 0.60 | 38.04 ± 3.92 | 6.83 ± 0.15 | 21.74 ± 1.84 | Significantly lower (p_adj = $1.93 \times 10^{-16}$); also significantly lower than TFI_5 (p_adj = $1.86 \times 10^{-4}$) |

**Note:** Note: The table corresponds to Fig. 4. Values are mean ± SD across n = 50 replicate simulations. Tumor burden comparisons were performed using the Mann–Whitney U test with Bonferroni correction across all 15 pairwise comparisons, with CONT as the reference.

**S2 Table.** Exhaustion state and recent cytotoxic activity of ACT products on Day 14 across TCE conditioning schedules, with comparisons against CONT.

| metric | group | U | p_raw | cliffs_delta | median_diff | ci_low | ci_high | p_adj |
|---|---|---|---|---|---|---|---|---|
| **Exhaustion state** | TFI_7 | 2500 | 7.04E-18 | 1 | 1.0175 | 0.9275 | 1.0875 | 3.52E-17 |
| **Exhaustion state** | TFI_5 | 2500 | 7.05E-18 | 1 | 0.67 | 0.60375 | 0.72625 | 3.52E-17 |
| **Exhaustion state** | TFI_3 | 942.5 | 0.034291 | -0.246 | -0.0375 | -0.11 | 0.017236 | 0.171453 |
| **Exhaustion state** | TFI_2 | 1842 | 4.54E-05 | 0.4736 | 0.13375 | 0.05125 | 0.201259 | 0.000227 |
| **Exhaustion state** | MO_FR | 1168 | 0.574168 | -0.0656 | 0 | -0.0675 | 0.055 | 1 |
| **Recent cytotoxic activity** | TFI_7 | 1061.5 | 0.19496 | -0.1508 | -0.7425 | -1.99 | 0.695063 | 0.974798 |
| **Recent cytotoxic activity** | TFI_5 | 172 | 1.10E-13 | -0.8624 | -4.2125 | -6.075 | -2.87744 | 5.51E-13 |
| **Recent cytotoxic activity** | TFI_3 | 1792 | 0.000189 | 0.4336 | 1.4775 | 0.3125 | 2.732563 | 0.000946 |
| **Recent cytotoxic activity** | TFI_2 | 1661.5 | 0.004606 | 0.3292 | 1.3425 | 0.07475 | 2.595 | 0.02303 |
| **Recent cytotoxic activity** | MO_FR | 1094.5 | 0.285272 | -0.1244 | -0.5675 | -1.85 | 0.6675 | 1 |

**Note:** The table corresponds to Fig. 4C. Comparisons were performed using the Mann–Whitney U test with Bonferroni correction within each metric; Cliff's δ and 95% confidence intervals are shown.

**S3 Table.** Pairwise comparisons of tumor burden on ACT Day 16 across TCE conditioning schedules.

| g1 | g2 | U | p_raw | cliffs_delta | median_diff | ci_low | ci_high | p_adj |
|---|---|---|---|---|---|---|---|---|
| **CONT** | **TFI_7** | 2490 | 1.28630498175 38502e-17 | 0.992 | 3280 | 2690.988 | 3787.013 | 1.92945747263 07752e-16 |
| **CONT** | **TFI_5** | 2424 | 5.97030034177 0663e-16 | 0.9392 | 1958.5 | 1518 | 2421.02499999 99996 | 8.95545051265 5994e-15 |
| **CONT** | **TFI_3** | 1017 | 0.10897167100 211709 | -0.1864 | -117 | -281 | 81.5124999999 9982 | 1 |
| **CONT** | **TFI_2** | 1324.5 | 0.60994722363 85701 | 0.0596 | 42.5 | -108 | 331.012499999 9998 | 1 |
| **CONT** | **MO_FR** | 929.5 | 0.027381 | -0.2564 | -190.5 | -307.513 | -6 | 0.41071565914 12184 |
| **TFI_7** | **TFI_5** | 615.5 | 1.23851740335 99414e-05 | -0.5076 | -1321.5 | -1969.54 | -601.5 | 0.000186 |
| **TFI_7** | **TFI_3** | 1 | 7.50236518089 5634e-18 | -0.9992 | -3397 | -3897.5 | -2768.96 | 1.12535477713 4345e-16 |
| **TFI_7** | **TFI_2** | 16.5 | 1.89335400471 33158e-17 | -0.9868 | -3237.5 | -3737.54 | -2593.46 | 2.84003100706 99737e-16 |
| **TFI_7** | **MO_FR** | 2 | 7.96702015069 0287e-18 | -0.9984 | -3470.5 | -3969.01 | -2842 | 1.19505302260 35431e-16 |
| **TFI_5** | **TFI_3** | 42 | 8.48195896738 522e-17 | -0.9664 | -2075.5 | -2543.51 | -1596.5 | 1.27229384510 7783e-15 |
| **TFI_5** | **TFI_2** | 111.5 | 4.32134408813 7166e-15 | -0.9108 | -1916 | -2380.01 | -1400 | 6.48201613220 5749e-14 |
| **TFI_5** | **MO_FR** | 37 | 6.33692578517 538e-17 | -0.9704 | -2149 | -2583 | -1676 | 9.50538867776 3069e-16 |
| **TFI_3** | **TFI_2** | 1586.5 | 0.02053909671 0802517 | 0.2692 | 159.5 | -45.5125 | 454 | 0.30808645066 203777 |
| **TFI_3** | **MO_FR** | 1185 | 0.65656978061 66135 | -0.052 | -73.5 | -253.013 | 136.012499999 99982 | 1 |
| **TFI_2** | **MO_FR** | 851.5 | 0.006073 | -0.3188 | -233 | -521 | -34 | 0.091099 |

**Note:** The table corresponds to Fig. 4E. Comparisons were performed using the Mann–Whitney U test with Bonferroni correction across all 15 pairwise comparisons; Cliff's δ and 95% confidence intervals are shown.

**S4 Table.** Exhaustion state and recent cytotoxic activity, and total CD8 T cell count of ACT products on Day 14 across schedules varying TFI timing and total TCE exposure.

| metric | g1 | g2 | U | p_raw | cliffs_delta | median_diff | ci_low | ci_high | p_adj |
|---|---|---|---|---|---|---|---|---|---|
| Exhaustion state | TFI_7 | TFI_7_offFirst | 2500 | 7.01E-18 | 1 | 0.81 | 0.7475 | 0.8875 | 4.21E-17 |
| Exhaustion state | TFI_7 | TFI_1on13off | 2500 | 7.04E-18 | 1 | 4.69 | 4.5975 | 4.78 | 4.22E-17 |
| Exhaustion state | TFI_7 | TFI_13off1on | 2500 | 5.46E-18 | 1 | 4.845 | 4.8025 | 4.92 | 3.27E-17 |
| Exhaustion state | TFI_7_offFirst | TFI_1on13off | 2500 | 7.00E-18 | 1 | 3.88 | 3.78 | 3.95 | 4.20E-17 |
| Exhaustion state | TFI_7_offFirst | TFI_13off1on | 2500 | 5.43E-18 | 1 | 4.035 | 3.99 | 4.085 | 3.26E-17 |
| Exhaustion state | TFI_1on13off | TFI_13off1on | 2160 | 3.08E-10 | 0.728 | 0.155 | 0.095 | 0.242562 | 1.85E-09 |
| Recent cytotoxic activity | TFI_7 | TFI_7_offFirst | 0 | 7.06E-18 | -1 | -47.1 | -48.225 | -45.4275 | 4.24E-17 |
| Recent cytotoxic activity | TFI_7 | TFI_1on13off | 2473.5 | 3.42E-17 | 0.9788 | 5.91 | 5.032437 | 6.755 | 2.05E-16 |
| Recent cytotoxic activity | TFI_7 | TFI_13off1on | 2500 | 7.03E-18 | 1 | 11.64 | 10.86744 | 12.3675 | 4.22E-17 |
| Recent cytotoxic activity | TFI_7_offFirst | TFI_1on13off | 2500 | 7.06E-18 | 1 | 53.01 | 51.38744 | 53.8075 | 4.24E-17 |
| Recent cytotoxic activity | TFI_7_offFirst | TFI_13off1on | 2500 | 7.03E-18 | 1 | 58.74 | 57.0925 | 59.425 | 4.22E-17 |
| Recent cytotoxic activity | TFI_1on13off | TFI_13off1on | 2500 | 7.03E-18 | 1 | 5.73 | 5.3375 | 6.2225 | 4.22E-17 |
| Total CD8 T Cells | TFI_7 | TFI_7_offFirst | 0 | 7.03E-18 | -1 | -1016.5 | -1033 | -1000.49 | 4.22E-17 |
| Total CD8 T Cells | TFI_7 | TFI_1on13off | 0 | 7.06E-18 | -1 | -6964.5 | -7114.51 | -6901.5 | 4.23E-17 |
| Total CD8 T Cells | TFI_7 | TFI_13off1on | 0 | 7.00E-18 | -1 | -2012.5 | -2026 | -2002 | 4.20E-17 |
| Total CD8 T Cells | TFI_7_offFirst | TFI_1on13off | 0 | 7.04E-18 | -1 | -5948 | -6096.51 | -5883.5 | 4.22E-17 |
| Total CD8 T Cells | TFI_7_offFirst | TFI_13off1on | 0 | 6.98E-18 | -1 | -996 | -1010 | -986 | 4.19E-17 |
| Total CD8 T Cells | TFI_1on13off | TFI_13off1on | 2500 | 7.01E-18 | 1 | 4952 | 4891 | 5088.075 | 4.20E-17 |

**Note:** The table corresponds to Fig. 5C. Comparisons were performed using the Mann–Whitney U test with Bonferroni correction across all six pairwise comparisons; Cliff's δ and 95% confidence intervals are shown.

**S5 Table.** Pairwise comparisons of tumor burden on ACT Day 16 across schedules varying TFI timing and total TCE exposure.

| g1 | g2 | U | p_raw | cliffs_delta | median_diff | ci_low | ci_high | p_adj |
|---|---|---|---|---|---|---|---|---|
| TFI_7 | TFI_7_offFirst | 1461.5 | 0.145779 | 0.1692 | 147 | -387.037 | 868.5625 | 0.874672 |
| TFI_7 | TFI_1on13off | 2500 | 7.07E-18 | 1 | 16861.5 | 16342 | 17453.5 | 4.24E-17 |
| TFI_7 | TFI_13off1on | 2500 | 7.07E-18 | 1 | 15591 | 15056 | 16182.5 | 4.24E-17 |
| TFI_7_offFirst | TFI_1on13off | 2500 | 7.06E-18 | 1 | 16714.5 | 16216.98 | 17067 | 4.24E-17 |
| TFI_7_offFirst | TFI_13off1on | 2500 | 7.06E-18 | 1 | 15444 | 14944.46 | 15809.01 | 4.24E-17 |
| TFI_1on13off | TFI_13off1on | 300.5 | 6.06E-11 | -0.7596 | -1270.5 | -1590.01 | -960 | 3.64E-10 |

**Note:** The table corresponds to Fig. 5E. Comparisons were performed using the Mann–Whitney U test with Bonferroni correction across all six pairwise comparisons; Cliff's δ and 95% confidence intervals are shown.
